# Coordinated immune, chloroplast and chemical defences underpin multilayered resistance to barley yellow dwarf virus and its aphid vector associated with the *Hordeum bulbosum*-derived Ryd4 introgression in barley

**DOI:** 10.64898/2026.07.28.741184

**Authors:** Amma L. Simon, Dasuni P. Jayaweera, Ilma A Qonaah, Daniel Verburg, Faith Clarke, Dong-Hyun Kim, Benjamin Urquhart, James P.E. Melichar, Tom C. Giles, Toby J.A. Bruce, Rumiana V. Ray

## Abstract

Barley yellow dwarf virus (BYDV), transmitted by the bird cherry-oat aphid (*Rhopalosiphum padi* L.), is among the most damaging viral diseases of barley, but the mechanisms underlying resistance to both the virus and its vector remain poorly understood. Here, we investigated resistance associated with the *Hordeum bulbosum*-derived Ryd4 introgression in the barley hybrid SY Kestrel by integrating behavioural, electrophysiological, physiological and multi-omics analyses with functional validation of defence metabolites.

SY Kestrel exhibited constitutive volatile-mediated antixenosis together with strong post-settlement antibiosis characterised by impaired phloem feeding, reduced aphid fitness and suppression of BYDV gene expression 10 days after transmission. Integrated transcriptomic, metabolomic and small RNA analyses revealed coordinated immune activation, chloroplast remodelling and defence metabolism associated with the resistance introgression. Candidate immune regulators were identified both within the refined Ryd4 interval and the surrounding introgressed region. Maintenance of photosystem II function was accompanied by reprogramming of α-linolenic acid-derived oxylipin metabolism, while phenylpropanoid and branched-chain amino acid/lysine pathways generated metabolites that directly reduced aphid survival.

We demonstrate that the Ryd4 introgression coordinates constitutive vector deterrence with host defence reprogramming to restrict aphid colonisation and suppress BYDV establishment. These results provide a mechanistic framework for improving durable resistance to aphid-transmitted viruses in cereals.

## Introduction

Successful transmission of vector-borne plant viruses depends on complex interactions between the insect vector, the pathogen and the host. However, the mechanisms that enable plants to resist both the vector and the virus remain largely unknown. The bird cherry-oat aphid (*Rhopalosiphum padi* L.) is the principal vector of barley yellow dwarf virus (BYDV), one of the most economically important viral pathogens of cereals worldwide. BYDV is transmitted in a persistent, circulative, non-propagative manner, with BYDV-PAV representing the dominant serotype in Europe (Gray & Gildow, 2003; Jiménez *et al*., 2020). Infection causes yellow dwarf disease, characterised by chlorosis, stunting and substantial yield losses (Jarošová *et al*., 2013; Ali *et al*., 2018). Although virus infection frequently disrupts photosynthetic performance, the contribution of chloroplasts to coordinated defence against both aphid vectors and viral pathogens is largely unexplored. Climate change, prolonged aphid migration and increasing insecticide resistance are expected to intensify disease pressure and reduce the effectiveness of chemical control, placing greater reliance on durable host resistance (Habekuß *et al*., 2009; Foster *et al*., 2014; Mc Namara *et al*., 2020). Plant resistance to aphids comprises antixenosis, which reduces host selection, and antibiosis, which impairs insect performance after feeding (Kogan & Ortman, 1978; Stout, 2013). Both mechanisms can reduce virus transmission by limiting vector settlement, probing and sustained phloem ingestion (Qonaah *et al*., 2026). In contrast, resistance to BYDV acts after inoculation by restricting virus establishment, replication or movement within the host (Cooper, 1983). Because successful virus transmission requires sequential completion of host selection, probing, sustained phloem feeding and viral establishment, resistance may arise through integrated defence layers acting across successive stages of infection rather than through independent mechanisms. Whether such defence programmes simultaneously restrict aphid colonisation and BYDV establishment remains unclear.

Several loci conferring resistance or tolerance to BYDV have been incorporated into barley breeding programmes, including Ryd1, Ryd2 and, more recently, Ryd4, introgressed from *Hordeum bulbosum* accession A17 (Suneson, 1955; Schaller *et al*., 1963; Habekuß *et al*., 2004). The Ryd4 introgression provides broad protection against BYDV-PAV, BYDV-MAV and the cereal yellow dwarf virus (CYDV)-RPV. Ryd4-mediated resistance was originally identified by the absence of detectable virus following aphid-mediated inoculation, with resistant plants remaining ELISA-negative after challenge with multiple virus species (Habekuß *et al*., 2004; Pidon *et al*., 2024). Subsequent fine mapping narrowed the resistance interval to a small region on chromosome 3H containing candidate nucleotide-binding leucine-rich repeat (NLR) and ankyrin-repeat genes, suggesting an immune receptor-mediated mechanism (Pidon *et al*., 2024). Although reduced aphid feeding was previously reported in the donor accession *Hordeum bulbosum* A17 (Schliephake *et al*., 2013), recombinant introgression lines carrying Ryd4 showed no evidence of altered aphid feeding behaviour, suggesting that resistance is mediated primarily through restriction of virus establishment rather than altered vector performance (Pidon *et al*., 2024). Whether the Ryd4 introgression deployed in elite commercial barley hybrids coordinates multilayered defence extending beyond antiviral immunity to influence aphid behaviour, host physiology and defence metabolism remains unknown.

Here, we investigated the mechanisms underlying resistance associated with the Ryd4 introgression in the commercial barley hybrid SY Kestrel by comparing it with the closely related susceptible hybrid Tektoo, which lacks the introgression. We hypothesised that resistance associated with the Ryd4 introgression extends beyond antiviral immunity by coordinating multiple defence layers that collectively restrict aphid colonisation and BYDV establishment. To test this hypothesis, we combined aphid behavioural assays, electrical penetration graph (EPG) analyses, chlorophyll fluorescence phenotyping and volatile profiling with transcriptomic, metabolomic and small RNA sequencing. Integrated multi-omics analyses were then used to identify coordinated defence networks and prioritise candidate resistance pathways for functional validation.

## Materials and methods

### Plant material, aphid cultures and growth conditions

Two closely related commercial hybrid barley (*Hordeum vulgare* L.) cultivars differing in the presence of the *Hordeum bulbosum*-derived Ryd4 resistance introgression were used throughout the study: SY Kestrel, which carries the Ryd4 resistance introgression together with additional linked *H. bulbosum* chromosomal segments, and Tektoo, which lacks the introgression. The two hybrids were generated using the same female parental line and highly similar male restorer lines, providing a closely matched genetic comparison for evaluating resistance associated with the Ryd4 introgression. Seed was provided by Syngenta Ltd. (Cambridgeshire, UK).

Plants were grown in Levington Advance Seed & Modular F2S compost (Evergreen Garden Care, UK) under controlled environment (20°C day,18°C night, 16h photoperiod). Seedlings were established in 8 × 12 modular trays (3.5L x 3.5W x 8D cm) before being transplanted into 1L pots at growth stage (GS) 10 (Zadoks *et al*., 1974) . Unless otherwise stated, experiments were performed using three-leaf stage plants (GS13) maintained at 22°C day, 18°C night under a 16 h photoperiod.

A parthenogenetic clone of the bird cherry-oat aphid (*Rhopalosiphum padi* L.) obtained from Rothamsted Research (Harpenden, UK) was maintained on oat (*Avena sativa* L.) cv. Gerald and used throughout the study. Viruliferous and non-viruliferous aphid populations were generated from this same clonal lineage to ensure that virus infection status, rather than aphid genotype, was the only experimental variable. Viruliferous aphids were produced by allowing aphids a four-day acquisition access period on BYDV-PAV-infected oat plants supplied by NIAB (Cambridge, UK), followed by transfer to healthy oat plants. Non-viruliferous aphids were maintained continuously on healthy oat plants. Virus infection status was routinely confirmed by reverse transcription quantitative PCR (RT-qPCR) using BYDV-specific primers (Balaji *et al*., 2003). Aphid colonies were maintained in BugDorm insect cages (47.5 cm³; NHBS Ltd., Devon, UK) at 20°C day, 16°C night, 60 ± 5% relative humidity and a 16 h photoperiod. Alate aphids were generated by colony overcrowding.

### Experimental workflow

The resistance phenotype associated with the Ryd4 introgression was investigated by integrating behavioural, electrophysiological, physiological and molecular analyses (Figs. 1,3). Whole-plant behavioural assays, volatile analyses, electrical penetration graph (EPG) recordings, chlorophyll fluorescence measurements and virus quantification were performed using the resistant and susceptible barley genotypes challenged with either non-viruliferous or viruliferous *R. padi*. Matched biological samples collected at 5 and 10 days post infestation (dpi) were subjected to transcriptomic, metabolomic and small RNA sequencing to enable direct multi-omics integration. Candidate defence pathways identified through supervised and unsupervised integrative analyses were subsequently validated using synthetic volatile compounds and artificial diet bioassays.

**Fig 1.**
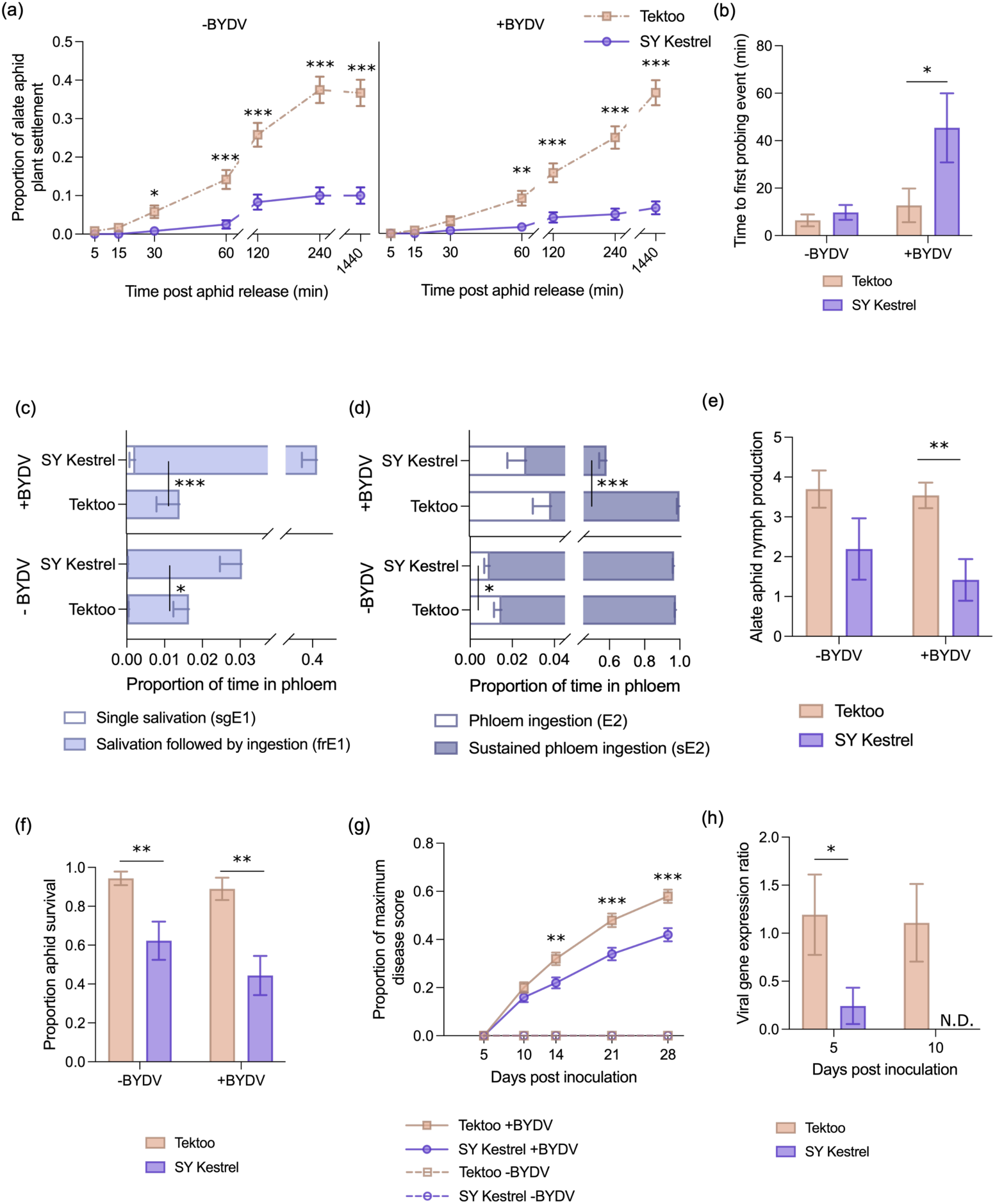
Constitutive and induced resistance associated with the Ryd4 introgression restricts aphid colonisation, phloem feeding and BYDV establishment in barley. (a) Settlement of alate non-viruliferous (-BYDV) and viruliferous (+BYDV) *R. padi* on the barley hybrids SY Kestrel (Ryd4 introgression) and Tektoo (lacking the introgression) at 5, 15, 30, 60, 120, 240 and 1440 min after aphid release (n = 6). Electrical penetration graph (EPG) analyses showing (b) time to first probe, (c) duration of single salivation (sgE1) and salivation preceding phloem ingestion (frE1) and (d) total (E2) and sustained (>10 min) phloem ingestion (sE2) (n = 10-13). Aphid performance measured as (e) alate fecundity and (f) apterous aphid survival (n = 5). BYDV infection assessed by (g) visual symptom scores (n = 10) and (h) relative viral gene expression (n = 3). Data are presented as back-transformed means ± SE. Statistical analyses were performed using generalised linear models (GLMs) with appropriate error distributions. Asterisks indicate significant differences between treatments (Fisher’s LSD, * *P* ≤ 0.05; ** *P* ≤ 0.01; *** *P* ≤ 0.001). N.D, no viral gene expression detected.

### Alate host selection and settlement behaviours

Host selection and settlement behaviour of viruliferous and non-viruliferous alate *R. padi* were assessed using whole-plant dual-choice assays. Experiments with viruliferous and non-viruliferous aphids were conducted independently in BugDorm cages partitioned into two choice chambers connected by a central release arena. One SY Kestrel and one Tektoo plant (9 cm pots) were positioned in opposing chambers, with genotype positions alternated between replicates to minimise positional bias.

Twenty alate aphids starved for 1 h were released into the central arena, and aphid settlement on each plant was recorded after 5, 15, 30, 60, 120, 240 and 1440 min. Cages were rotated by 90° every 60 min during the first 240 min to minimise directional effects. Nymph production was recorded after 24 h. Each treatment comprised six biological replicates and the entire experiment was repeated independently.

### Volatile Collection and GC-MS Analysis

Volatile organic compounds (VOCs) were collected from Tektoo and SY Kestrel plants under three treatments: uninfested controls, infestation with non-viruliferous aphids and infestation with viruliferous aphids. Headspace collections were performed at 2 and 7 dpi using dynamic air entrainment adapted from Birkett *et al*., (2004) and Qonaah *et al*., (2024). Plants receiving aphid treatments were infested with ten first-instar aphids per plant and maintained individually within whole-plant cages between collections. Volatiles were trapped on Porapak Q filters over 24 h, eluted with 750 μl dichloromethane and stored at -20°C prior to analysis. Three biological collections were performed for each treatment.

VOC profiling was carried out using an Agilent 7820A gas chromatograph coupled to a 5977B mass selective detector (Agilent Technologies, UK) following Ali *et al*., (2022). Samples were analysed in a single randomised analytical batch together with pooled quality-control samples, solvent blanks and n-alkane standards. Compound identities were assigned by comparison with mass spectral libraries and confirmed by co-chromatography with authentic standards (100 ng μl⁻¹) as described by Asamoah *et al*., (2025).

### Olfactometer bioassays

Behavioural responses of viruliferous and non-viruliferous alate *R. padi* to plant VOCs were assessed using four-arm olfactometers (Pettersson, 1970) as described by Qonaah *et al*., (2024). Separate assays were conducted for each barley genotype and sampling time point (2 and 7 dpi). Olfactometer arms contained VOC collections from uninfested plants, plants infested with non-viruliferous aphids, plants infested with viruliferous aphids and blank controls.

Individual aphids were observed for 16 min and movement recorded using the olfactometerR package in R (v4.1.2). To minimise directional bias, the olfactometer was rotated by 45° every 2 min. Ten biological replicates were completed for each treatment under controlled environmental conditions (25 °C, 28 ±2% humidity).

Behavioural responses to individual synthetic VOCs were evaluated separately using one treatment arm containing the test compound and three solvent control arms (diethyl ether). Synthetic compounds were tested at concentrations equivalent to those quantified in plant headspace collections.

### Volatile priming effects on aphid fecundity

To determine whether perception of host-derived VOCs influenced subsequent aphid performance, viruliferous and non-viruliferous alate aphids were pre-exposed for 1 h to VOCs collected from uninfested Tektoo or SY Kestrel plants, or to blank solvent controls, following Asamoah *et al*., (2025).

Following exposure, two alate aphids were confined to the first leaf of a Tektoo seedling (GS19) using clip cages. Nymph production was recorded after 24 and 48 h, and aphid survival was assessed after 48 h. Mean daily fecundity was calculated as the average number of nymphs produced per surviving aphid per day. Each treatment comprised six biological replicates and was repeated independently.

To assess the contribution of individual VOCs, aphids were similarly pre-exposed to synthetic compounds at biologically relevant concentrations before transfer to Tektoo seedlings. Survival and fecundity were quantified after 48 h. Compounds enriched in Tektoo and SY Kestrel volatile blends were evaluated in separate experiments comprising five biological replicates repeated independently.

### Aphid antibiosis assay

Antibiosis was evaluated by measuring survival and reproductive performance on Tektoo and SY Kestrel. Viruliferous and non-viruliferous aphids were assessed simultaneously in separate insect cages to prevent cross-contamination. A single alate aphid was confined to the first leaf of each plant using a clip cage. Survival was assessed after 23 h before removing any surviving adults and retaining one randomly selected apterous aphid from each replicate for further 7 days when total nymph production was recorded. Offspring survival was subsequently assessed after an additional seven days. All surviving aphids developed into apterous morphs during the experiment. Two independent experiments with six biological replicates per treatment were conducted under controlled-environment conditions.

### Electrical Penetration Graph (EPG) analysis

Feeding behaviour of third- and fourth-instar viruliferous and non-viruliferous *R. padi* on Tektoo and SY Kestrel were monitored using Giga-8dd Direct current (DC) electrical penetration graph (EPG) system (EPG systems, Wageningen, The Netherlands) (Tjallingii & Hogen Esch, 1993) following Qonaah *et al*., (2024). Aphids were tethered with gold wire and placed individually on the first leaf of each plant in a 9cm pot. Recordings of viruliferous and non-viruliferous aphids were conducted independently under constant illumination at 23°C and 28 ± 2% relative humidity. Recordings were excluded if aphids failed to produce waveforms in the first hour or exhibited more than three hours of continuous inactivity. At least 11 valid biological replicates were obtained per treatment (Table S1). Waveforms were annotated manually and feeding parameters calculated using the EPG-Excel analysis macro.

### Chlorophyll fluorescence

To investigate whether coordinated defence associated with the Ryd4 introgression extends to chloroplast function during aphid infestation, Tektoo and SY Kestrel plants were analysed using chlorophyll a fluorescence induction kinetics (OJIP transients). Plants were positioned in 8 × 12 seed trays in a completely randomised design within 47.5cm^3^ BugDorm cages. Plants were arranged in a completely randomised design, with aphid treatments maintained in separate insect cages to prevent cross-contamination. Ten second-instar aphids were confined to the first leaf of each plant using clip cages for 2 d before removal. Control plants received empty clip cages. Six biological replicates were included for each genotype and treatment.

OJIP measurements were performed every 2 d from 2-10 dpi using a FluorPen FP100 fluorometer (Photon Systems Instruments, Czech Republic) on the youngest fully expanded leaf between 10:00 and 15:00 h (Ajigboye *et al*., 2017). Fluorescence transients were induced using a saturating light pulse of 3000 μmol m⁻² s⁻¹. Parameters calculated included absorption flux per reaction centre (ABS/RC), trapping flux per reaction centre (TRo/RC), electron transport flux per reaction centre (ETo/RC), dissipated energy flux per reaction centre (DIo/RC), quantum yield (QY) and Fv/Fm’.

### Sampling for transcriptomic, metabolomic and small RNA analyses

To characterise molecular responses associated with aphid infestation and BYDV infection, Tektoo and SY Kestrel plants were assigned to three treatments: uninfested controls, infestation with non-viruliferous aphids and infestation with viruliferous aphids. Plants were arranged in a randomised design with aphid treatments maintained in separate BugDorm cages.

Ten first-instar aphids were confined to the first leaf of each plant using clip cages. Aphid survival was assessed at 5 and 10 dpi, and nymph production quantified at 10 dpi. The first and second leaves were harvested at 5 and 10 dpi (n = 5 biological replicates), immediately frozen in liquid nitrogen and stored at -80°C. Matched biological samples were subsequently used for transcriptomic, metabolomic and small RNA sequencing to enable direct multi-omics integration.

### Untargeted metabolomic profiling

Freeze-dried leaf tissue (50mg) was extracted in pre-cooled methanol:water (80:20, v/v) containing 0.1% formic acid at a final concentration of 10mg ml^-1^. Samples were homogenised on a rotary shaker (300rpm, 1hour) on ice and centrifuged at 14,000rpm for 10 min at 4°C. Supernatants were removed and centrifuged again before storage at -80°C prior to analysis.

Metabolomic profiling was performed using Q-Exactive Plus Orbitrap mass spectrometer (MS) coupled to a Dionex U3000 UHPLC system (Thermo Fisher Scientific, Hertfordshire, UK). Metabolites (10 µl, 4°C) were separated via a ZIC-*p*HILIC column (4.6 × 150 mm, 5μm particle size, Merck SeQuant, Watford, UK) maintained at 45 °C. The mobile phase was initially set at 20% A (20 mM ammonium carbonate in water) and 80% B (acetonitrile), followed by a linear increase to 95% A over 8 min. Then the gradient was then returned to the starting conditions over 2 min, and subsequently the column was left to re-equilibrate for 4 mins and flowrate restored for 1min. The total runtime was 15min. The MS was operated in ESI+ and ESI− switching acquisition modes for LC-MS profiling of the samples and in data-dependent MS/MS (ddMS/MS) for identification for the analysis of the quality control (QC) samples. For MS parameters, spray voltage was 4.5 kV (ESI+) and 3.5 (ESI−), and capillary voltage was 20 V (ESI+) and −15 V (ESI−). The sheath, auxiliary and sweep gas flow rates were 40, 5 and 1 (arbitrary unit), respectively, for both modes. Capillary and heater temperatures were maintained at 275 and 150 °C, respectively. Data were acquired for the LC-MS profiling with a resolution of 70 000 from *m*/*z* 70–1050. Top 5 ddMS/MS was performed on the QC sample (*n* = 3) at a resolution of 17 500 and a stepped normalised collision energy (NEC) of 20, 30 and 40.

Samples were randomised and analysed in a single analytical batch together with pooled QC samples, reagent blanks and authentic standards. Pooled QC (*n* = 8) were injected at the beginning of the analysis to condition the column and after every 6 samples to assess the robustness and repeatability of the analytical system. Raw data was processed and metabolite annotation using performed Compound Discoverer (version 3.3 SP1) through accurate mass matching, retention time alignment and spectral comparisons with authentic standards and public databases including BioCyc, Metabolika and ChemSpider. Mass tolerance for peak picking and metabolite identification was ≤ 5 ppm for fragment ions, minimum peak intensity was 1.0 × 10^6^, intensity tolerance was at 30%, area max was greater or equal to 3.0 × 10^6^, maximum retention time shift was 0.25 min and activation energy tolerance was 5. Identified metabolites were assigned as confidence level 2 or 3, based on the recommendation by the Chemical Analysis Working Group, Metabolomics Standards Initiative (MSI) (Sumner *et al*., 2007).

### Transcriptome sequencing and differential expression analysis

Total RNA was extracted using RNeasy Plant kit (Qiagen) whilst RNase-free DNase kit (Qiagen) was used to remove residual genomic DNA. RNA integrity and concentration were assessed prior to mRNA sequencing by BMK GENE (Cambridge, UK) using the Illumina NovaSeq X/DNBSEQ-T7 platform.

Raw reads were quality filtered to remove adapter sequences and low-quality sequences as described by Burchardt *et al*., (2024). Clean reads were aligned to the barley reference genome assembly GCF_904849725.1 (<u>NCBI</u>). Differential gene expression analyses were performed between barley genotypes, aphid treatments and sampling time points using standard RNA-seq workflows. Differentially expressed genes were identified following multiple-testing correction using a false discovery rate (FDR) threshold of <0.05.

### Small RNA sequencing and miRNA analysis

Small RNA libraries were prepared from the same RNA samples used for transcriptome sequencing (n = 5 biological replicates per treatment and sampling time point) using standard library construction method with cDNA purification performed with AmpureXP beads, according to the manufacturer’s instructions. Libraries were sequenced on a high-throughput PE150 sequencing platform to generate single-end 18-30 bp reads.

Raw reads were processed by removing adaptor sequences, low-quality reads and sequences outside the expected small RNA size range using an in-house perl script. Filtered reads were mapped to the barley reference genome (GCF_904849725.1) using bowtie2 and BLAST. Known miRNAs were identified by comparison with miRBase v22, while novel miRNAs were predicted using miRDeep2 software (Friedländer *et al*., 2012) based on characteristic precursor secondary structures.

Read counts were normalised using transcript per kilobase (TPM) algorithm and differential expression analyses were performed using edgeR (Robinson *et al*., 2010). Differentially expressed miRNAs were identified using an FDR-adjusted *P* < 0.05. Putative target genes were predicted using TargetFinder (Bo & Wang, 2005) and integrated with transcriptomic data to identify negatively correlated miRNA-mRNA pairs associated with resistance responses.

### Metabolite validation assays

Candidate metabolites prioritised through multi-omics analyses were functionally evaluated using artificial diet bioassays. Artificial diets were supplemented with 3-O-feruloyl-D-quinic acid, isobutyric acid, L-serine or saccharopine at concentrations of 0.01, 0.05 or 0.25 mg ml⁻¹ following De Zutter *et al*., (2016). Viruliferous and non-viruliferous aphids were assessed simultaneously under controlled-environment conditions.

Three second- or third-instar aphids were introduced into each feeding chamber (n = 5), and survival recorded after 3 and 5 days. Diet consumption was estimated gravimetrically by measuring sachet weight before and after feeding. Median effective concentrations (EC₅₀) were calculated using Abbott-corrected mortality as described by Drakulic *et al.,* (2016).

### Multi-omics integration

Transcriptomic, metabolomic and small RNA datasets generated from matched biological samples were integrated to identify coordinated molecular signatures associated with resistance to viruliferous and non-viruliferous *R. padi*. For differential expression analysis, transcript and small RNA counts were analysed using count-based statistical models, with normalisation factors and dispersion estimates derived from the raw count data using DESeq2 (Love *et al*., 2014). Variance-stabilised or transformed abundance values were used only for exploratory visualisation and integration, not as the input for differential expression testing. Prior to MOFA integration, RNA abundance values, metabolite abundances and small RNA abundances were log-transformed where appropriate and autoscaled so that features from each omics layer contributed on a comparable scale.

Two complementary analytical approaches were employed. First, supervised analyses, including differential expression analysis and orthogonal partial least squares-discriminant analysis (OPLS-DA), were used to identify genes, metabolites and miRNAs discriminating barley genotypes and aphid treatments. Secondly, unsupervised Multi-Omics Factor Analysis (MOFA2) (Argelaguet *et al*., 2020) was used to identify latent factors representing the principal sources of variation shared across molecular datasets without predefined sample classes. Transcriptomic, metabolomic and small RNA datasets were incorporated as independent data views within a single MOFA model. Candidate molecular features consistently identified by both supervised analyses and MOFA-derived latent factors were functionally annotated using KEGG pathways and Gene Ontology classifications before prioritisation for experimental validation.

### RT-qPCR validation of candidate transcripts

Expression of selected candidate genes was validated using reverse transcription quantitative PCR (RT-qPCR). Complementary DNA was synthesised from RNA samples (n = 3 biological replicates) using the iScript cDNA synthesis kit (Bio-Rad). RT-qPCR was performed using SYBR Green chemistry on a CFX96 Touch Real-Time PCR Detection System (Bio-Rad). Amplification conditions consisted of an initial denaturation at 95°C for 10 min followed by 40 amplification cycles of 95°C for 15 s, 60°C for 1 min and 72°C for 10 s. Primer sequences are listed in Table S2. Relative transcript abundance was calculated using 2^−ΔΔC^ method (Livak & Schmittgen, 2001) following normalisation against the reference genes *GAPDH* and *TUBB* (Jarošová & Kundu, 2010).

### BYDV symptom assessment and viral quantification

Visual BYDV symptom development was assessed using a modified 0-5 disease severity scale adapted from (Choudhury *et al*., 2019), where 0 represented asymptomatic plants and 5 represented >60% leaf chlorosis. Symptom assessments were performed at 7, 14 and 21 dpi for most experiments, with additional evaluations at 28 dpi for EPG experiments and at 5, 10, 14, 21 and 28 dpi for omics experiments.

BYDV accumulation was quantified by RT-qPCR using RNA extracted from transcriptomic/metabolomic samples harvested at 5 and 10 dpi (n = 3). cDNA synthesis and amplification were performed as described above using BYDV-specific primers (Balaji *et al*., 2003; Qonaah *et al*., 2026). Relative viral transcript abundance of BYDV-PAV coat protein was normalised against *GAPDH* using the 2⁻ΔΔCt method (Jarošová *et al*., 2013)

### Statistical analysis

All statistical analysis was performed using GenStat® 24^th^ Edition (VSN International Ltd, Hemel Hempstead, UK) unless otherwise stated. Generalised linear models (GLMs) were fitted using distributions appropriate for each response variable. Count data were analysed using Poisson distributions with log link functions, proportional data using binomial distributions with logit link functions, and continuous variables assuming Gaussian errors. Model assumptions were assessed by inspection of residual plots and dispersion statistics where appropriate. Statistical significance was accepted at *P* < 0.05.

Alate host selection, aphid performance, chlorophyll fluorescence and RT-qPCR data were analysed using GLMs followed by appropriate post hoc comparisons. Olfactometer responses to synthetic volatile compounds were analysed using Mann-Whitney U-tests. Viruliferous and non-viruliferous aphid datasets were analysed independently for EPG experiments.

Volatile and metabolomic datasets were analysed using SIMCA (v13; Sartorius Stedim Data Analytics, Sweden). Data were transformed and scaled before orthogonal partial least squares-discriminant analysis (OPLS-DA). Significant features were identified using variable importance in projection (VIP > 1) together with adjusted P < 0.05. Model robustness was assessed using R², Q² and permutation testing following (Eriksson *et al*., 2006) with full statistics provided in Tables S3, S4 and S5.

## Results

### The Ryd4 resistance introgression combines constitutive antixenosis, post-settlement antibiosis and suppression of BYDV establishment

Whole-plant choice assays demonstrated that both non-viruliferous and viruliferous *R. padi* preferentially settled on the susceptible genotype Tektoo (-Ryd4), with significant differences becoming apparent after 30 min and 60 min following release, respectively (Fig. **1a**). Settlement on SY Kestrel (+Ryd4) remained consistently lower throughout the experiment, demonstrating constitutive antixenosis against both aphid populations. Aphid settlement on SY Kestrel was more than 4 times lower than on Tektoo.

Electrical penetration graph (EPG) analysis revealed that resistance extended beyond host acceptance to disrupt compatible phloem feeding. Viruliferous aphids required significantly longer to initiate probing on SY Kestrel than on Tektoo (Fig. **1b**), while both aphid populations spent less total time probing on the resistant genotype (Fig. **S1a**). Aphids reached the sieve elements in both genotypes, with no significant difference in initial phloem access (Fig. **S1b**). Following phloem contact, however, feeding behaviour diverged. Aphids feeding on SY Kestrel exhibited prolonged salivation within sieve elements (Fig. **1c**), particularly when viruliferous, together with a pronounced reduction in sustained phloem sap ingestion (sE2) (Fig. **1d**). Additional EPG variables showed increased stylet derailment for viruliferous aphids and reduced pathway activity on SY Kestrel for non-viruliferous aphids (Fig. **S1b,c**), consistent with impaired establishment of compatible sieve element feeding rather than delayed phloem location. Reduced host acceptance and feeding resulted in lower aphid fitness. Alate fecundity was significantly reduced on SY Kestrel, with viruliferous aphids producing approximately 40% fewer nymphs than on Tektoo (Fig. **1e**). Apterous aphids exhibited reduced survival, lower reproductive output and decreased second-generation nymph survival on the resistant genotype irrespective of virus status (Fig. **1f**; Fig. **S1d,e**).

Resistance also extended beyond vector performance to suppression of BYDV establishment. Plants carrying the Ryd4 introgression developed significantly fewer visual disease symptoms than Tektoo (Fig. **1g**), while viral transcript abundance was markedly reduced and became undetectable by 10 dpi (Fig. **1h**).

### Constitutive volatile blends mediate aphid host discrimination and influence aphid fitness

To determine the basis of constitutive antixenosis, volatile organic compounds (VOCs) emitted by Tektoo and SY Kestrel were analysed following aphid feeding and BYDV infection. Olfactometer assays demonstrated that host discrimination was strongly mediated by volatile cues. Both non-viruliferous and viruliferous alates were consistently attracted to volatile blends emitted by Tektoo but avoided those from SY Kestrel (Fig. **2a**, Fig. **S2a**).

**Fig 2.**
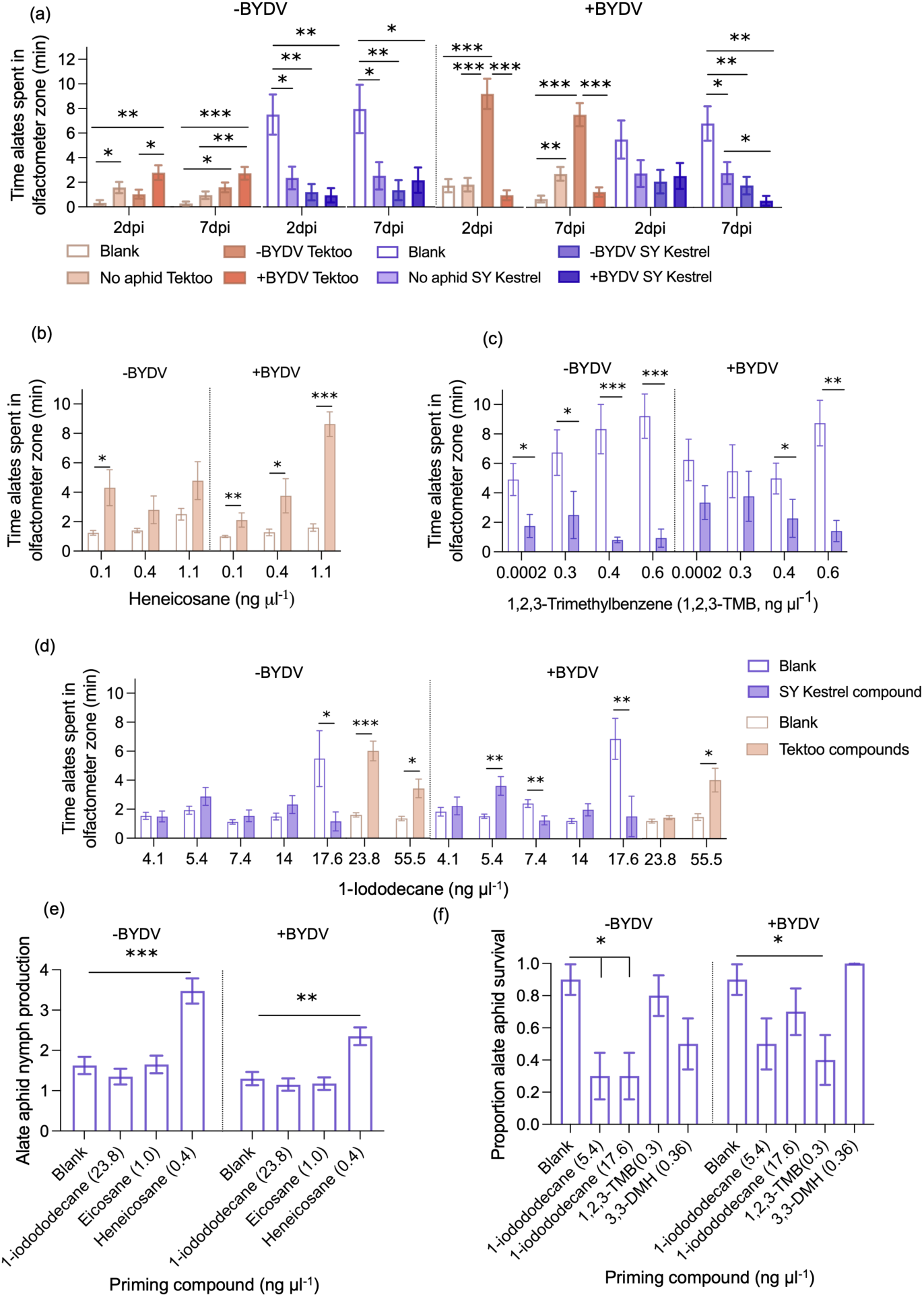
A constitutive volatile blend associated with the Ryd4 introgression deters aphid host selection and reduces aphid fitness. **(a)** Behavioural responses of alate viruliferous (+BYDV) and non-viruliferous (-BYDV) *R. padi* to volatile organic compounds (VOCs) collected from the barley hybrids SY Kestrel (Ryd4 introgression) and Tektoo (lacking the introgression) Olfactometer assays compared responses to blank controls and headspace VOCs from uninfested plants and plants infested with non-viruliferous or viruliferous aphids at 2 and 7 d post infestation (dpi) (n = 10). Behavioural responses to synthetic VOCs identified by GC-MS: **(b)** heneicosane, **(c)** 1,2,3-trimethylbenzene and **(d)** 1-iodododecane (n = 10). Aphid performance following 1 h volatile priming showing **(e)** nymph production and **(f)** survival (n = 5). Data are presented as back-transformed means ± SE. Statistical analyses were performed using generalised linear models (GLMs) or Mann–Whitney U-tests, as appropriate. Asterisks indicate significant differences between treatments. * *P* ≤ 0.05; ** *P* ≤ 0.01; *** *P* ≤ 0.001.

Although aphid feeding and BYDV infection modified volatile profiles in both genotypes (Tables **S3**, **S4**), behavioural consequences differed substantially. Non-viruliferous aphids preferentially responded to VOCs emitted from BYDV-infected Tektoo plants, whereas viruliferous aphids were more strongly attracted to VOCs from healthy Tektoo plants (Fig. **2a**; Fig. **S2a**). In contrast, volatile blends from SY Kestrel repelled both aphid populations at both 2 and 7 dpi irrespective of infestation status (Fig. **2a**; Fig. **S2a**), indicating that constitutive chemical differences in volatile composition dominated host recognition.

Comparative GC-MS profiling identified five discriminatory semiochemicals between the two genotypes (Table **S6, S7, Fig. S2b,c**). Heneicosane and eicosane were detected exclusively in Tektoo volatile blends, whereas 1,2,3-trimethylbenzene and 3,3-dimethylhexane were specific to SY Kestrel. Olfactometer assays with authentic standards revealed contrasting behavioural responses. Heneicosane elicited strong attraction responses in both aphid populations, whereas 3,3-dimethylhexane and 1,2,3-trimethylbenzene induced concentration-dependent repellence, particularly in viruliferous aphids (Fig. **2b**,**c**; Fig. **S2d,e**). Responses to 1-iodododecane varied with concentration and aphid infection status, suggesting that quantitative differences in this compound contribute to host discrimination (Fig. **2d**).

Perception of host-derived volatile blends altered aphid performance independently of feeding. Priming non-viruliferous alates with SY Kestrel VOCs significantly reduced survival and fecundity compared with aphids exposed to Tektoo VOCs or blank controls (Fig. **S2f,g**). Among individual compounds, heneicosane increased nymph production in both aphid populations, whereas 1-iodododecane and 1,2,3-trimethylbenzene reduced survival and reproductive output (Fig. **2e,f**).

### The Ryd4 resistance introgression differentially regulates PSII photochemistry during viral challenge

To determine whether *Ryd4*-mediated resistance was accompanied by altered photosynthetic performance, chlorophyll fluorescence parameters were monitored from 2 to 10 dpi (Fig. **3**, Fig. **S3**). Aphid feeding altered PSII activity in both genotypes, however the physiological responses differed according to host genotype and aphid infection status.

**Fig 3.**
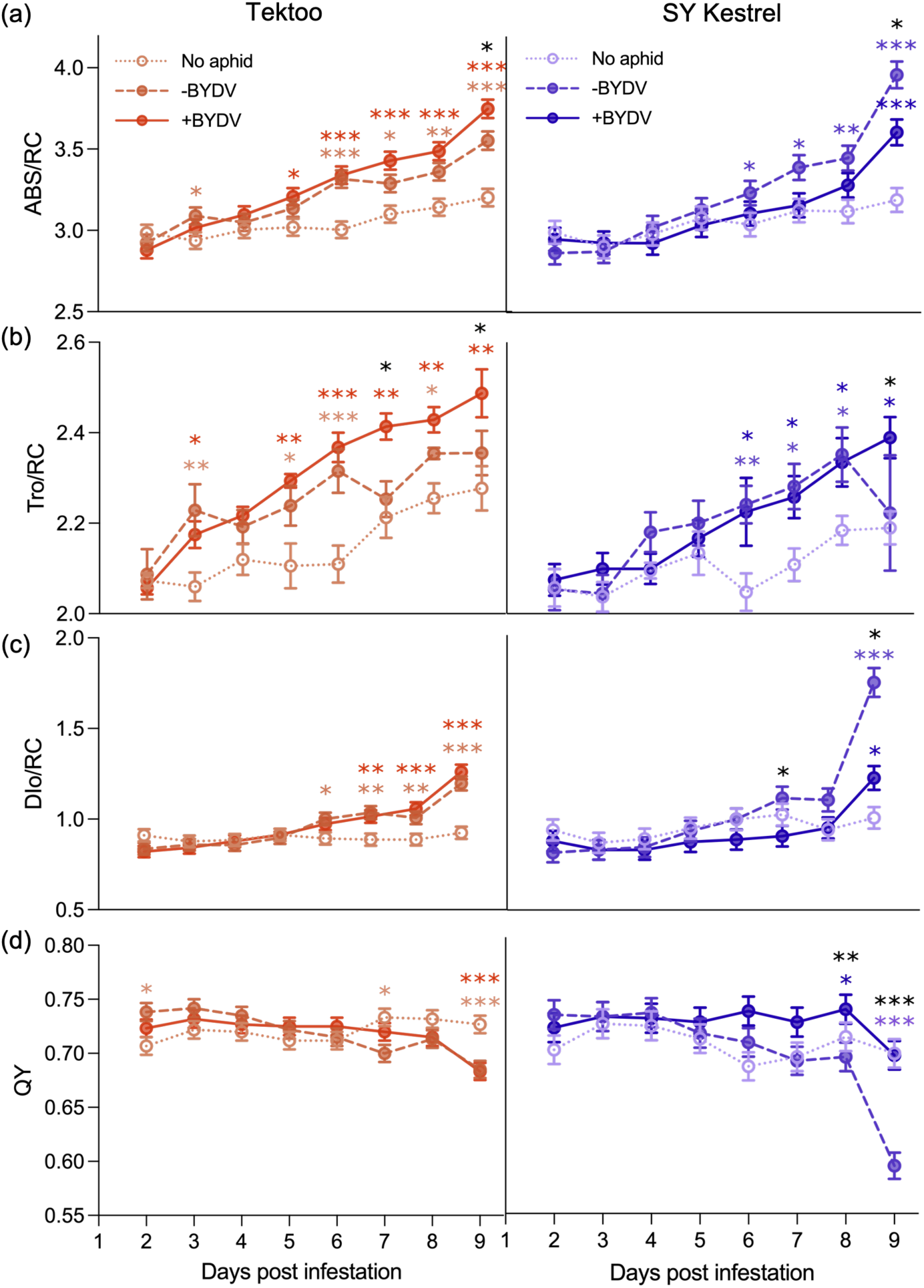
The Ryd4 introgression preserves photosynthetic energy fluxes during aphid infestation. Chlorophyll fluorescence parameters measured in the barley hybrids SY Kestrel (Ryd4 introgression) and Tektoo (lacking the introgression) following infestation with non-viruliferous (-BYDV) or viruliferous (+BYDV) *R. padi*, or in uninfested controls, from 2 to 9 d post infestation (dpi). **(a)** absorption per reaction centre (ABS/RC), (**b)** trapping energy flux per reaction centre (TRo/RC), (**c)** dissipated energy flux per reaction centre (DIo/RC) and (**d)** PSII quantum yield (Qy) (*n* = 6). Data are presented as back-transformed means ± SE. Statistical analyses were performed using generalised linear models (GLMs). Black asterisks indicate significant differences between -BYDV and +BYDV treatments, whereas coloured asterisks indicate significant differences relative to the corresponding uninfested control. * *P* ≤ 0.05; ** *P* ≤ 0.01; *** *P* ≤ 0.001.

In Tektoo, infestation by both non-viruliferous and viruliferous aphids increased energy absorption (ABS/RC), trapping (TRo/RC), electron transport (ETo/RC) and energy dissipation (DIo/RC) per reaction centre from 3-5 dpi onwards (Fig. **3a-c**, Fig. **S3a**). These changes culminated in significant reductions in quantum yield (QY) and PSII efficiency (Fv/Fm’) by 9 dpi (Fig. **3d**; Fig. **S3b**).

In contrast to Tektoo, SY Kestrel displayed divergent responses depending on aphid infection status. Feeding by non-viruliferous aphids increased energy dissipation and reduced trapping efficiency, resulting in lower QY and Fv/Fm′ than healthy controls (Fig. **3**, Fig. **S3**). In contrast, during viruliferous aphid infestation, electron transport, trapping efficiency and energy absorption progressively recovered while QY and Fv/Fm′ remained comparable with non-infested plants (Fig. **3**, **S3**).

### Integrated multi-omics identify three defence modules associated with the Ryd4 resistance introgression

To investigate the molecular basis of resistance associated with the Ryd4 resistance introgression, transcriptomic, metabolomic and small RNA datasets generated from matched biological samples were analysed using complementary supervised and unsupervised integration approaches. Initial metabolomic analyses separated resistant and susceptible genotypes across aphid treatments in OPLS-DA models (Fig. **S4a,b**), while differential abundance analyses identified metabolites distinguishing the two genotypes (Tables **S5, S8**). Integration of these features with transcriptomic and small RNA datasets, together with unsupervised Multi-Omics Factor Analysis (MOFA) separated resistant and susceptible plants according to genotype and aphid treatment (Fig. **S4**).

Comparison of differentially expressed genes with the recently refined 66.5 kb Ryd4 interval identified candidate genes within the introgression together with immune-associated genes immediately flanking the mapped region (Table **S9**). Constitutively upregulated *Ryd4_CNL2* mapped to the same genomic position as the complete *CC-NLR* proposed by Pidon et al. (2024), whereas *ANK*, the second candidate gene within the interval, showed higher expression in the susceptible genotype Tektoo (Fig. **4a**,**b**). Among flanking immune-associated genes, *WAK* and *Annexin* exhibited constitutively higher expression in SY Kestrel, *PIK6-NP* was induced at both sampling times following infestation by either aphid population, and *BAK1* was strongly induced at 10 dpi with viruliferous aphids (Fig. **4b**).

**Fig 4.**
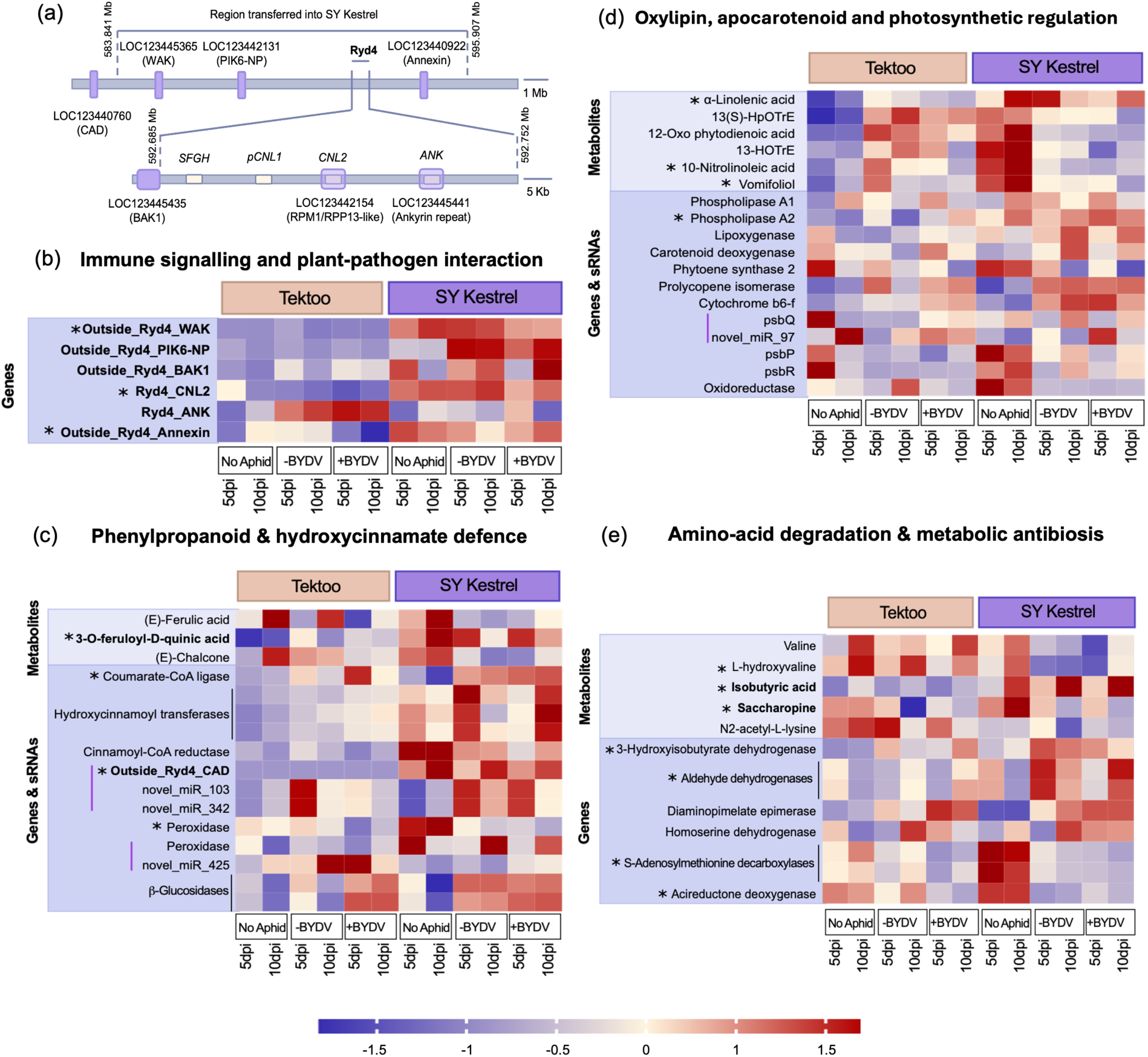
Integrated multi-omics identifies coordinated immune, chloroplast and metabolic defence modules associated with the Ryd4 introgression. **(a)** Genomic organisation of the Ryd4 introgression showing candidate genes previously identified within the refined Ryd4 interval (white) and differentially expressed genes identified in SY Kestrel within and flanking the introgressed region (purple). Curated defence modules identified through concordance between supervised multi-omics integration, unsupervised multi-omics factor analysis (MOFA) and functional enrichment analyses showing **(b)** immune and plant-pathogen interaction genes associated with the introgression, **(c)** phenylpropanoid and hydroxycinnamate metabolism chloroplast-associated oxylipin, **(d)** apocarotenoid and photosynthetic regulation, and **(e)** branched-chain amino acid and lysine metabolism associated with metabolic defence. Features validated experimentally are shown in **bold**, MOFA-supported features are indicated by asterisks (*), and lines adjacent to small RNA features denote their predicted target genes.

To identify downstream defence mechanisms, molecular features consistently prioritised by both integration approaches were combined with Gene Ontology (GO) and KEGG enrichment analyses. This framework converged on three robust defence modules (Fig. **4c-e**).

The first integrated defence module comprised phenylpropanoid metabolism, characterised by accumulation of hydroxycinnamate-derived metabolites, including ferulic acid, 3-O-feruloyl-D-quinic acid and chalcone, together with increased expression of *4CL*, *HCT*, *CCR*, *CAD*, peroxidases and β-glucosidases (Fig. **4c**). Differential expression of regulatory small RNAs targeting *CAD* and peroxidase transcripts suggested additional post-transcriptional regulation of this pathway.

The second module linked chloroplast-associated oxylipin metabolism with photosynthetic function. Under non-infested conditions, SY Kestrel contained higher basal levels of 13(S)-HpOTrE, OPDA, 13-HOTrE, 10-nitrolinoleic acid and vomifoliol than Tektoo (Fig **4d**). Aphid infestation reversed this pattern, with these metabolites declining in SY Kestrel but increasing in Tektoo. In contrast, the oxylipin precursor α-linolenic acid remained consistently more abundant in SY Kestrel irrespective of aphid treatment (Fig **4d**).

These metabolic changes were accompanied by differential expression of phospholipases, lipoxygenase, carotenoid biosynthesis enzymes and multiple photosystem II-associated genes, including *PsbP*, *PsbQ*, *PsbR*, cytochrome *b6f* components and chloroplast oxidoreductases (Fig. **4d**). Reciprocal expression of *PsbQ* and the predicted regulatory *novel_miR_97* was observed exclusively in the resistant genotype.

The third module comprised branched-chain amino acid and lysine metabolism. SY Kestrel accumulated higher levels of L-hydroxyvaline, isobutyric acid and saccharopine, accompanied by altered expression of genes involved in branched-chain amino acid catabolism, lysine biosynthesis and methionine recycling, including *HIBADH*, *ALDH*, *DapF*, *HSDH*, *SAMDC* and *ARD* (Fig **4e**).

Independent enrichment analyses supported these curated defence modules. GO analysis identified enrichment of defence responses, photosynthetic function, hormone signalling and amino acid metabolism (Fig. **S5**), while KEGG analysis identified enrichment of phenylpropanoid biosynthesis, plant-pathogen interaction, carotenoid biosynthesis, plant hormone signalling and amino acid metabolism (Fig. **S6**).

To validate metabolites prioritised by the integrated multi-omics analyses, representative compounds from the phenylpropanoid and branched-chain amino acid/lysine metabolism modules were tested in artificial feeding assays at 0.01, 0.05 and 0.25 ng μl⁻¹ (Fig. **5**). Simplified pathway reconstructions incorporating differentially expressed genes illustrate the position of each metabolite within its respective pathway (Fig. **5a**,**c**).

**Fig 5.**
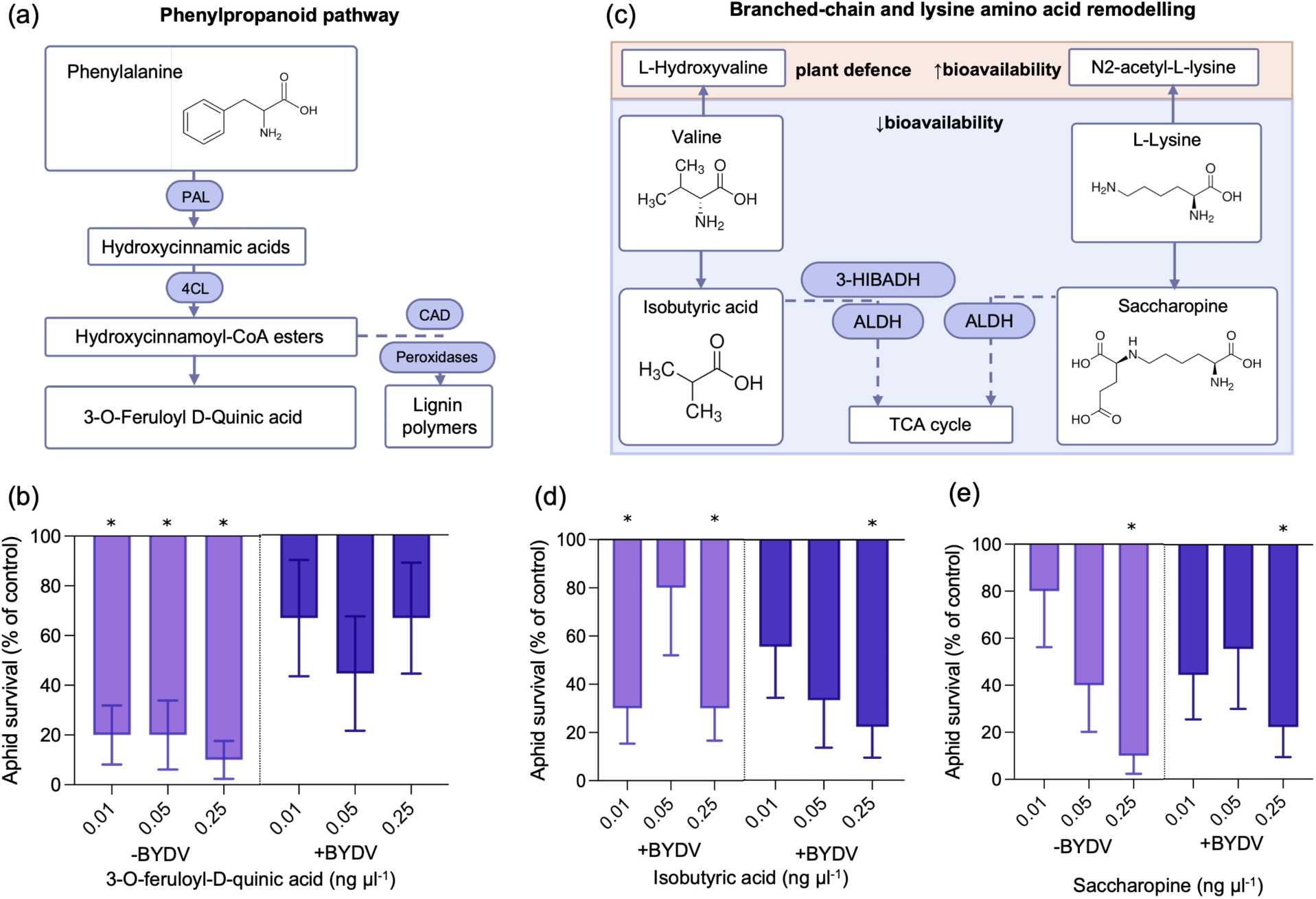
Functional validation confirms aphicidal activity of defence metabolites identified by integrated multi-omics. **(a)** Simplified phenylpropanoid pathway highlighting the accumulation of 3-O-feruloyl-D-quinic acid in SY Kestrel. **(b)** Effects of 3-O-feruloyl-D-quinic acid on *R. padi* survival at 0.01, 0.05, 0.25 ng µl^-1^ in artificial diet assays (n = 5). **(c)** Simplified branched-chain amino acid and lysine metabolic pathways highlighting isobutyric acid and saccharopine, which accumulated in SY Kestrel. Effects of **(d)** isobutyric acid and **(e)** saccharopine on aphid survival at 0.01, 0.05, 0.25 ng µl^-1^ in artificial diet assays (n = 5). Data are presented as back-transformed means ± SE. Aphid survival is expressed relative to the control artificial diet. Statistical analyses were performed using generalised linear models (GLMs). Asterisks indicate significant differences from the control. * *P* ≤ 0.05.

Within the phenylpropanoid module, 3-O-feruloyl-D-quinic acid reduced survival of non-viruliferous *R. padi* in a concentration-dependent manner but had no significant effect on viruliferous aphids (Fig. **5b**). Isobutyric acid significantly reduced survival of non-viruliferous aphids at 0.01 and 0.25 ng μl⁻¹, whereas viruliferous aphids were affected only at the highest concentration (Fig. **5d**). Similarly, saccharopine produced a concentration-dependent reduction in survival of non-viruliferous aphids, while both aphid populations were affected at 0.25 ng μl⁻¹ (Fig. **5e**).

## Discussion

Here, we show that resistance associated with the Ryd4 introgression extends beyond antiviral defence to comprise a coordinated, multilayered defence system acting before, during and after aphid feeding. Because SY Kestrel carries the Ryd4 introgression together with additional chromosomal segments derived from *H. bulbosum*, these mechanisms should be considered properties of the complete introgressed region rather than the refined Ryd4 interval alone. SY Kestrel combines constitutive volatile-mediated antixenosis, post-settlement antibiosis and suppression of BYDV establishment, thereby disrupting successive stages required for successful virus transmission. Collectively, these findings redefine resistance associated with the Ryd4 introgression as a coordinated, multilayered defence programme that establishes sequential barriers to virus transmission through interconnected immune, physiological and metabolic networks rather than acting solely through antiviral resistance.

Resistance to insect-transmitted viruses can arise through direct restriction of virus multiplication or indirectly through traits that reduce vector host selection and feeding (Qonaah *et al*., 2026). Our results provide strong evidence for the latter while also supporting an additional contribution of post-inoculation antiviral resistance associated with the Ryd4 introgression. Although reduced aphid settlement and sustained phloem feeding are expected to decrease virus inoculation, the coordinated activation of immune signalling, chloroplast function and defence metabolism, together with the marked suppression of viral accumulation, suggests that post-inoculation resistance mechanisms also contribute to limiting BYDV establishment. Prior to host contact, aphids consistently avoided SY Kestrel irrespective of their viruliferous status or plant infection history, demonstrating constitutive volatile-mediated antixenosis. This contrasted with the susceptible hybrid Tektoo, where viral infection modified volatile blends to induce the characteristic host-preference reversal previously described for BYDV-vector interactions, in which non-viruliferous aphids preferentially colonise infected hosts whereas viruliferous aphids select healthy plants (Ingwell *et al*., 2012; Hu *et al*., 2022). The absence of this behavioural switch in SY Kestrel indicates that constitutive host-derived volatile cues override virus-induced manipulation of vector behaviour.

Several volatile compounds were associated with this resistance phenotype. Heneicosane, previously linked with *R. padi* attraction in wheat (Asamoah *et al*., 2025), was detected only in Tektoo and increased aphid fecundity, supporting its role as a susceptibility-associated semiochemical. In contrast, 3,3-dimethylhexane and 1,2,3-trimethylbenzene were specific to SY Kestrel and elicited strong repellent responses, while 1-iodododecane reduced aphid fitness in a concentration-dependent manner. Although 1,2,3-trimethylbenzene has previously been implicated in insect behavioural responses in other plant systems (Arora *et al*., 2021; Odermatt *et al*., 2025), its role in cereal defence has not previously been reported. Likewise, this is, to our knowledge, the first study identifying 3,3-dimethylhexane as a barley volatile associated with aphid repellence. Together, these compounds define a constitutive volatile signature that disrupts vector host selection before probing begins and therefore establishes the first barrier to successful virus transmission.

Following aphid settlement, resistance shifted from behavioural deterrence to interference with compatible sieve element feeding. Viruliferous aphids exhibited delayed probing, increased stylet derailment, prolonged salivation and markedly reduced sustained phloem ingestion, resulting in reduced survival, lower fecundity and diminished BYDV accumulation. Similar reductions in phloem feeding have previously been reported in barley carrying *H. bulbosum*-derived resistance (Schliephake *et al*., 2013), suggesting that disruption of compatible sieve element feeding is a conserved feature of resistance associated with the introgression. Increased stylet derailment has frequently been associated with altered cell wall architecture and activation of cell wall-associated defence (Tjallingii, 1988), consistent with constitutive activation of *WAK* and *Annexin* together with inducible expression of *PIK6-NP* and *BAK1* in SY Kestrel. Together, these observations indicate that behavioural deterrence is reinforced by disruption of compatible phloem feeding, establishing a second barrier to successful virus transmission. Maintenance of chloroplast function and defence metabolism subsequently reinforce this resistance by restricting virus establishment following inoculation.

Consistent with this post-inoculation phase of resistance, one of the most striking features of the resistant phenotype was extensive remodelling of chloroplast function. Whereas susceptible Tektoo exhibited progressive reductions in PSII efficiency following aphid infestation, SY Kestrel maintained photochemical performance during viruliferous aphid challenge despite adopting a more photoprotective physiological state during feeding by non-viruliferous aphids. Although aphid feeding and virus infection are each known to perturb photosynthesis, our results show that resistance associated with the Ryd4 introgression is accompanied by coordinated maintenance of photosystem function together with extensive chloroplast transcriptional and metabolic reprogramming, placing chloroplasts at the centre of immune coordination during aphid-transmitted virus infection. Chloroplasts are increasingly recognised as central hubs of plant immunity integrating reactive oxygen species (ROS) production, lipid-derived signalling and phytohormone biosynthesis (de Torres Zabala *et al*., 2015; Serrano *et al*., 2016). Consistent with this role, integrated multi-omics analyses identified coordinated regulation of phospholipases, lipoxygenases, carotenoid biosynthesis genes and multiple photosystem II-associated proteins, including PsbP, PsbQ and PsbR, together with genotype-specific reciprocal expression of *PsbQ* and novel_miR_97. These observations are particularly noteworthy because the oxygen-evolving complex of PSII has emerged as a conserved target of numerous positive-sense RNA viruses, whose proteins directly interact with PsbP and PsbQ to suppress chloroplast ROS signalling and antiviral defence (Sui *et al*., 2006; Zhao *et al*., 2016; Bhattacharyya & Chakraborty, 2018; Li *et al*., 2026). Although comparable interactions have not yet been demonstrated for BYDV, the reciprocal regulation of *PsbQ* and novel_miR_97, together with the maintenance of PSII photochemical performance in SY Kestrel, raises the possibility that preservation of oxygen-evolving complex integrity sustains chloroplast-mediated immune signalling and contributes to restricting BYDV establishment. Elucidating whether this represents a conserved antiviral mechanism in luteovirus resistance will require future functional investigation.

Plastid remodelling extended beyond maintenance of photosynthetic performance to extensive reprogramming of lipid metabolism. SY Kestrel exhibited constitutively elevated α-linolenic acid together with contrasting accumulation of the α-linolenic acid-derived oxylipins OPDA, 13(S)-HpOTrE and 13-HOTrE following aphid infestation. Because these oxylipins are central regulators of defence signalling and stress acclimation (Wasternack & Feussner, 2018), these observations suggest that preservation of photosynthetic performance in SY Kestrel reflects active immune regulation rather than simply reduced virus-induced damage. Together, these findings indicate that maintenance of photosystem integrity and oxylipin signalling form an integrated plastid-centred defence programme contributing to restriction of BYDV establishment.

Beyond plastid-associated processes, integrated multi-omics identified two additional defence modules encompassing phenylpropanoid metabolism and branched-chain amino acid and lysine metabolism. Functional validation confirmed that representative metabolites from both pathways directly reduced aphid fitness, providing experimental support for predictions generated through systems-level integration and distinguishing these pathways from purely correlative multi-omics signatures. Among these metabolites, 3-O-feruloyl-D-quinic acid links enhanced phenylpropanoid metabolism with hydroxycinnamate-derived defence, whereas isobutyric acid and saccharopine reveal previously unrecognised roles for branched-chain amino acid and lysine metabolism in cereal resistance to aphids. Although phenylpropanoid-derived metabolites are well established contributors to structural and chemical defence (Dixon & Paiva, 1995), and isobutyric acid possesses recognised antimicrobial activity (Oka *et al*., 2015), their direct effects on cereal aphids have received little attention. These findings therefore establish a mechanistic link between systems-level predictions and biologically relevant defence functions while identifying candidate metabolites for future resistance breeding.

Within the recently refined Ryd4 interval identified by (Pidon *et al*., 2024), *Ryd4_CNL2* co-localised with the complete *CC-NLR* previously proposed as the leading candidate gene, whereas *ANK* was more highly expressed in susceptible Tektoo. Although functional validation will ultimately be required, these findings further strengthen *Ryd4_CNL2* as the leading candidate gene while also suggesting that additional immune regulators within the surrounding *H. bulbosum* introgression are likely to contribute to the coordinated multilayered defence programme described here.

## Conclusion

Successful transmission of aphid-borne viruses depends on the sequential completion of host selection, probing, sustained phloem feeding and viral establishment. Our results demonstrate that resistance associated with the Ryd4 introgression disrupts multiple stages within this transmission pathway through constitutive volatile-mediated repellence, interference with compatible phloem feeding, suppression of BYDV establishment and maintenance of host physiological function. Rather than relying solely on direct antiviral resistance, our findings support a model in which the Ryd4 introgression coordinates immune activation, chloroplast remodelling and defence metabolism together with vector-mediated resistance to limit BYDV establishment. By disrupting multiple barriers within the transmission process, resistance associated with the Ryd4 introgression is likely to provide greater durability than mechanisms acting at a single stage, while preservation of photosynthetic performance during viruliferous aphid challenge may help sustain crop productivity under disease pressure. More broadly, this work demonstrates that durable resistance to aphid-transmitted viruses can emerge through the coordinated disruption of multiple stages in the transmission process, providing a mechanistic framework for breeding resilient cereal crops.

## Acknowledgements

We would like to thank Miss Emma Vines for generating preliminary settlement data, Mr. John Clews for assistance in GC-MS, Dr. Kostya Kanyuka for providing +BYDV *R. padi* starting population and Dr Sandra Martinez-Jarquin for assistance in metabolite identification.

## Author contributions

RVR, BU and JM conceived the study. ALS, DHK, BU, JM, TJAB and RVR designed the experiments. ALS, DPJ, IAQ, DV and FC performed experimental work. TCG conducted multi-omics factor analysis (MOFA). DHK, TJAB and RVR supervised data analysis and interpretation. ALS and RVR wrote the original manuscript. All authors contributed to manuscript revision and approved the final version.

## Data Availability

Supplementary figures and tables are provided with this preprint. Scripts, code and processed datasets required to reproduce the Multi-Omics Factor Analysis (MOFA) will be made publicly available in a GitHub repository upon publication. Raw sequencing data will be deposited in an appropriate public repository before publication.

## Supporting Information

**Table S1.** Number of experiments and valid replicates included in electrical penetration graph (EPG) analyses of *Rhopalosiphum padi* feeding behaviour on the barley (*Hordeum vulgare*) hybrids SY Kestrel and Tektoo.

**Table S2**. Primers used to validate genes within and near region transferred into SY Kestrel in quantitative PCR.

**Table S3**. Model fitness (R^2^) and predictive ability (Q^2^) values of Orthogonal partial least squares-discrimination analysis (OPLS-DA) for comparisons between plant volatile profiles of barley (*Hordeum vulgare*) genotypes SY Kestrel and Tektoo under No aphid, non-viruliferous *Rhopalosiphum padi* infestation (-BYDV) and viruliferous *R. padi* infestation (+BYDV) collected at 2, 7 and 14-days post infestation (dpi).

**Table S4.** Model fitness (R^2^) and predictive ability (Q^2^) values of Orthogonal partial least squares-discrimination analysis (OPLS-DA) for comparisons between plant volatile profiles of No aphid, non-viruliferous *Rhopalosiphum padi* infestation (-BYDV) and viruliferous *R. padi* infestation (+BYDV) barley (*Hordeum vulgare*) genotypes SY Kestrel and Tektoo under collected at 2, 7 and 14-days post infestation (dpi).

**Table S5.** Model fitness (R^2^) and predictive ability (Q^2^) values of Orthogonal partial least squares-discrimination analysis (OPLS-DA) for comparisons between metabolome of barley (*Hordeum vulgare*) genotypes SY Kestrel and Tektoo under No aphid, non-viruliferous *Rhopalosiphum padi* infestation (-BYDV) and viruliferous *R. padi* infestation (+BYDV) 5 and 10-days post infestation (dpi).

**Table S6.** Relative abundances and statistical analyses of volatile organic compounds (VOCs) identified in barley volatile blends with SY Kestrel against Tektoo comparison.

**Table S7**. Relative abundances and statistical analyses of volatile organic compounds (VOCs) identified in barley volatile blends with -BYDV against +BYDV comparison.

**Table S8.** Variable importance in projection (VIP) scores from orthogonal partial least squares-discriminant analysis (OPLS-DA) and adjusted *P*-values for metabolites distinguishing the barley (*Hordeum vulgare*) hybrids SY Kestrel and Tektoo.

**Table S9**. Candidate genes within and flanking the refined Ryd4 introgression identified by transcriptomic and Multi-Omics Factor Analysis (MOFA) and validated by quantitative PCR.

**Figure S1.** Feeding behaviour and life-history traits of apterous non-viruliferous (-BYDV) and viruliferous (+BYDV) *Rhopalosiphum padi* on the barley (*Hordeum vulgare*) hybrids SY Kestrel and Tektoo.

**Figure S2**. Orthogonal partial least squares-discrimination analysis (OPLS-DA) plot of barley volatile organic compounds (VOCs) and behavioural responses of non-viruliferous (-BYDV) and viruliferous (+BYDV) alate *Rhopalosiphum padi* to barley chemical cues and its effects on aphid life-history traits

**Figure S3.** Chlorophyll fluorescence parameters of the barley (*Hordeum vulgare*) hybrids SY Kestrel and Tektoo following infestation with non-viruliferous (-BYDV) or viruliferous (+BYDV) *Rhopalosiphum padi*.

**Figure S4.** Orthogonal partial least squares-discrimination analysis (OPLS-DA) plot of barley metabolic profle and multi-omics factorial analysis (MOFA) of *Hordeum vulgare* with *Ryd4* gene (SY Kestrel) and its parent without *Ryd4* (Tektoo) treated with no aphids, non-viruliferous (-BYDV) and viruliferous (+BYDV) aphids.

**Figure S5.** Gene Ontology (GO) enrichment analysis of transcriptomic responses in the barley (*Hordeum vulgare*) hybrids SY Kestrel and Tektoo following infestation with non-viruliferous (-BYDV) or viruliferous (+BYDV) *Rhopalosiphum padi*.

**Figure S6.** Kyoto Encyclopedia of Genes and Genomes (KEGG) pathway enrichment analysis of transcriptomic responses in the barley (*Hordeum vulgare*) hybrids SY Kestrel and Tektoo following infestation with non-viruliferous (-BYDV) or viruliferous (+BYDV) *Rhopalosiphum padi*.

