## Supporting information for "Coordinated immune, chloroplast and chemical defences underpin multilayered resistance to barley yellow dwarf virus and its aphid vector associated with the *Hordeum bulbosum*-derived Ryd4 introgression in barley"

The following Supporting Information is available for this article:

**Table S1**. Number of trials and valid replicates conducted in electrical penetration graph (EPG) experiments on barley (*Hordeum vulgare*) SY Kestrel and Tektoo. Experiments using non-viruliferous (-BYDV) and viruliferous (+BYDV) *Rhopalosiphum padi* were conducted separately.

| Aphid | Genotype | Number of trials | Number of valid replicates |
|---|---|---|---|
| -BYDV | Tektoo | 14 | 11 |
| | SY Kestrel | 14 | 12 |
| +BYDV | Tektoo | 18 | 12 |
| | SY Kestrel | 17 | 12 |

**Table S2**. Primers used to validate genes within and near regions transferred into SY Kestrel in quantitative PCR.

| Gene | Gene name | Forward primer | Reverse primer |
|---|---|---|---|
| LOC123440760 | CAD | CCACACAATAATCGCGGCTG | TATCACGGACCGGAAAGCAC |
| LOC123445365 | WAK | TCCGGGTAAAGGCCAGAAAC | ATCGCCTTTCATTCCGGGTT |
| LOC123442131 | PIK6-NP | TATGGCCCAGCTACAGGGAT | AAGCCTCCAGCAAGTCTGTC |
| LOC123445435 | BAK1 | TACTCGCAAAGGCAGGTGTG | GAGTTGTCGAGGTTGCAGGT |
| LOC123442154 | CNL2 | GAGGCCTTGGTAGGAACAGC | TGAGGACGGCATGCATCAAT |
| LOC123445441 | ANK | GACGGCCTTCGTATTGGCTA | TGTGTCGCCATGCGTATTCT |
| LOC123440922 | Annexin | CCGCTGAGATGATCCGACAG | GCTCCCAACCGATCCATCAC |

**Table S3**. Model fitness ($R^2$) and predictive ability ($Q^2$) values of Orthogonal partial least squares-discrimination analysis (OPLS-DA) for comparisons between plant volatile profiles of barley (*Hordeum vulgare*) genotypes SY Kestrel and Tektoo under No aphid, non-viruliferous *Rhopalosiphum padi* infestation (-BYDV) and viruliferous *R. padi* infestation (+BYDV) collected at 2, 7 and 14-days post infestation (dpi).

| **Volatiles Collection Timepoint** | **Interaction** | **$R^2$X(cum)** | **$R^2$Y(cum)** | **$Q^2$(cum)** |
|---|---|---|---|---|
| 2dpi | SY Kestrel No aphid v Tektoo No aphid | 0.918 | 1 | 0.993 |
| | SY Kestrel –BYDV v Tektoo –BYDV | 0.565 | 0.998 | 0.976 |
| | SY Kestrel +BYDV v Tektoo +BYDV | 1 | 1 | 1 |
| 7dpi | SY Kestrel No aphid v Tektoo No aphid | 1 | 1 | 1 |
| | SY Kestrel –BYDV v Tektoo –BYDV | 1 | 1 | 1 |
| | SY Kestrel +BYDV v Tektoo +BYDV | 0.938 | 1 | 0.991 |
| 14dpi | SY Kestrel No aphid v Tektoo No aphid | 0.968 | 1 | 0.995 |
| | SY Kestrel –BYDV v Tektoo –BYDV | 1 | 1 | 1 |
| | SY Kestrel +BYDV v Tektoo +BYDV | 1 | 1 | 1 |

**Table S4.** Model fitness ($R^2$) and predictive ability ($Q^2$) values of Orthogonal partial least squares-discrimination analysis (OPLS-DA) for comparisons between plant volatile profiles of No aphid, non-viruliferous *Rhopalosiphum padi* infestation (-BYDV) and viruliferous *R. padi* infestation (+BYDV) barley (*Hordeum vulgare*) genotypes SY Kestrel and Tektoo under collected at 2, 7 and 14-days post infestation (dpi).

| Genotype | Volatile Collection Timepoint | Interaction | $R^2X$(cum) | $R^2Y$(cum) | $Q^2$(cum) |
|---|---|---|---|---|---|
| | | No aphid v -BYDV | 0.971 | 0.999 | 0.991 |
| | 2dpi | No aphid v +BYDV | 1 | 1 | 1 |
| | | –BYDV v +BYDV | 1 | 1 | 1 |
| | | No aphid v -BYDV | 0.957 | 1 | 0.994 |
| Tektoo | 7dpi | No aphid v +BYDV | 1 | 1 | 1 |
| | | –BYDV v +BYDV | 1 | 1 | 1 |
| | | No aphid v -BYDV | 0.991 | 0.999 | 0.993 |
| | 14dpi | No aphid v +BYDV | 1 | 1 | 1 |
| | | –BYDV v +BYDV | 0.489 | 0.397 | 0.0385 |
| | | No aphid v -BYDV | 1 | 1 | 1 |
| | 2dpi | No aphid v +BYDV | 1 | 1 | 1 |
| | | –BYDV v +BYDV | 1 | 1 | 1 |
| | | No aphid v -BYDV | 1 | 1 | 1 |
| SY Kestrel | 7dpi | No aphid v +BYDV | 1 | 1 | 1 |
| | | –BYDV v +BYDV | 0.892 | 1 | 0.993 |
| | | No aphid v -BYDV | 0.776 | 1 | 0.991 |
| | 14dpi | No aphid v +BYDV | 0.835 | 1 | 0.998 |
| | | –BYDV v +BYDV | 1 | 1 | 1 |

**Table S5.** Model fitness ($R^2$) and predictive ability ($Q^2$) values of Orthogonal partial least squares-discrimination analysis (OPLS-DA) for comparisons between metabolome of barley (*Hordeum vulgare*) genotypes SY Kestrel and Tektoo under No aphid, non-viruliferous *Rhopalosiphum padi* infestation (-BYDV) and viruliferous *R. padi* infestation (+BYDV) 5 and 10-days post infestation (dpi).

| Aphid treatment | Interaction | $R^2X$(cum) | $R^2Y$(cum) | $Q^2$(cum) |
|---|---|---|---|---|
| | SY Kestrel v Tektoo 5d | 0.274 | 0.661 | 0.0394 |
| No aphid | SY Kestrel v Tektoo 10d | 0.817 | 1 | 0.851 |
| | SY Kestrel v Tektoo 5d | 0.581 | 0.986 | 0.801 |
| -BYDV | SY Kestrel v Tektoo 10d | 0.807 | 0.994 | 0.864 |
| | SY Kestrel v Tektoo 5d | 0.883 | 1 | 0.836 |
| +BYDV | SY Kestrel v Tektoo 10d | 0.884 | 1 | 0.953 |

### Figure S1

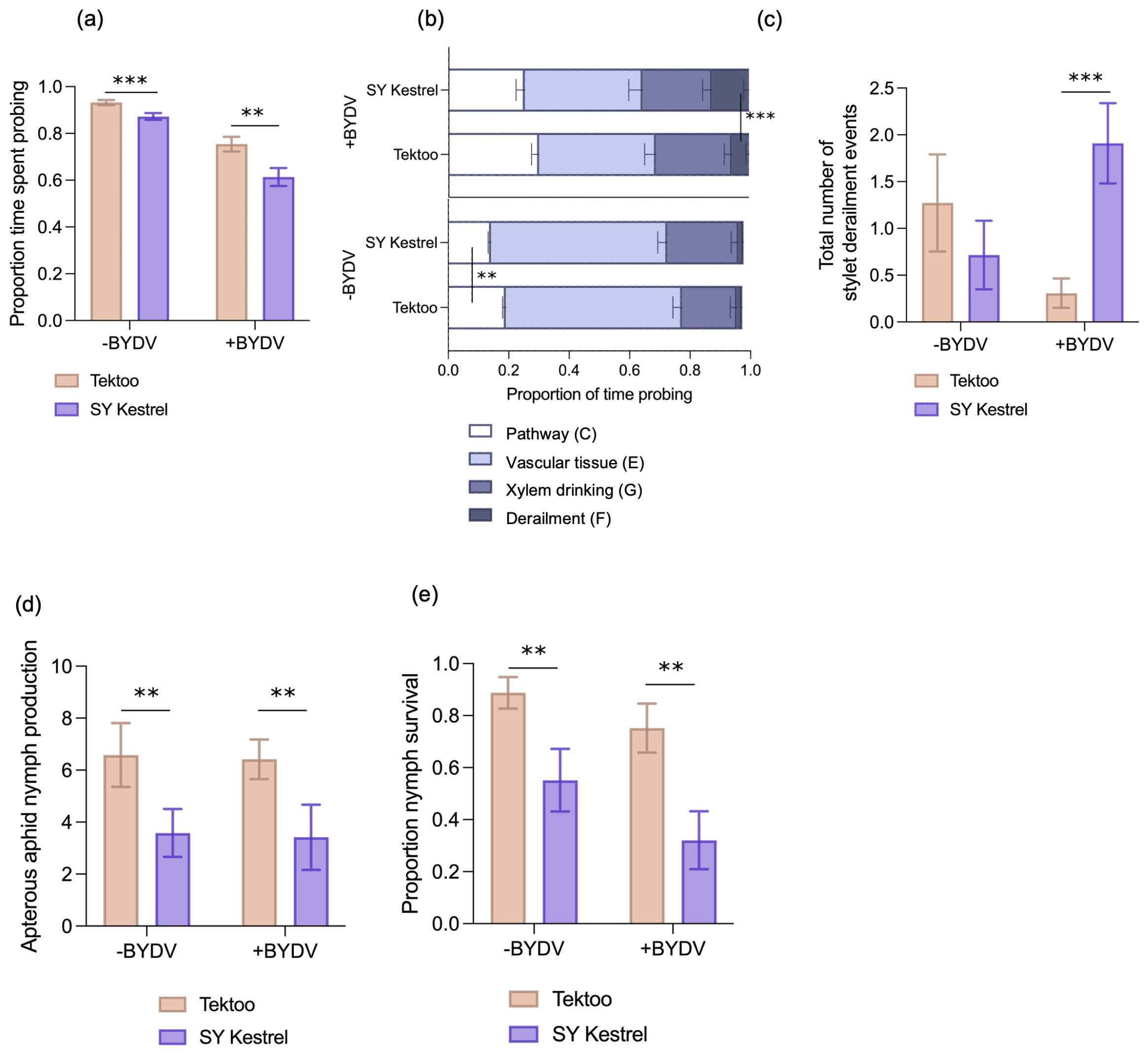


**Fig S1.** Apterous non-viruliferous (-BYDV) and viruliferous (+BYDV) *Rhopalosipum padi* feeding behaviour, fecundity and survival on the barley hybrids SY Kestrel (Ryd4 introgression) and Tektoo (lacking the introgression). Electrical penetration graph (EPG) waveforms indicate **(a)** Proportion of time spent probing, **(b)** proportion of time aphid spent in different leaf tissues, and **(c)** total number of stylet derailment events (n = 10-13). **(d)** Number of nymphs produced by apterous aphids (n = 6). **(e)** proportion of nymph survival (n = 6). Proportion of feeding events and survival were analysed using generalised linear model (GLM) with binomial distribution and link function logit. Number of stylet derailment and nymph production were analysed using GLM with poisson distribution and link function log. Back-transformed means shown, Fisher's least significant difference (LSD) applied for pairwise comparison indicated by asterisk * $P \leq 0.05$; ** $P \leq 0.01$; *** $P \leq 0.001$

### Figure S2

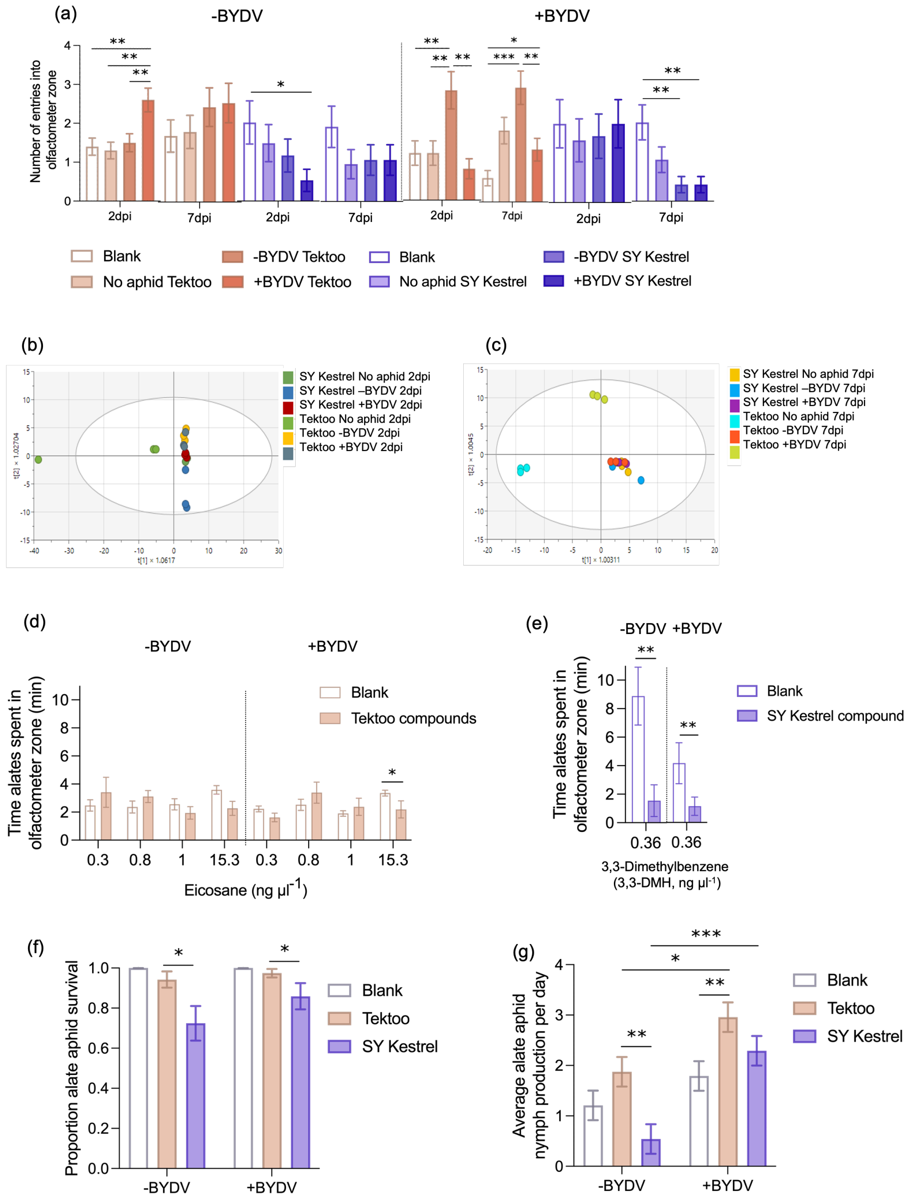


**Fig S2**. Behavioural responses of non-viruliferous (-BYDV) and viruliferous (+BYDV) alate *Rhopalosiphum padi* to barley volatile organic compounds (VOCs). **(a)** Number of entries into olfactometer containing blank, headspace VOC of plants treated with no aphids, non-virulifeorus aphids, and viruliferous aphids collected at 2- and 7dpi (n = 10). Orthogonal partial least squares-discrimination analysis (OPLS-DA) plot of barley VOCs and VOCs at **(b) 2**dpi and **(c) 7**dpi. Behavioural responses to synthetic VOCs identified by GC-MS: Time alates spent in olfactometer containing **(d)** eicosane identified in Tektoo VOCs, **(e)** 3,3-dimethylbenzene identified in SY Kestrel VOCs (n = 10). Aphid life-history traits following 1 h priming with VOCs from SY Kestrel and Tektoo **(f)** alate survival and **(g)** average nymph production by alate (n = 5). Olfactometer assay with plant headspace VOCs and nymph production were analysed using generalised linear model (GLM) with poisson distribution and link function log. Aphid survival was analysed using GLM with binomial distribution and link function logit. Olfactometer assays with synthetic compounds were analysed with non-parametric Mann-Whitney U test.

Asterisk indicate significance from Fisher's least significant difference (LSD) in GLM analysis and Mann-Whitney U test * $P \leq 0.05$; ** $P \leq 0.01$; *** $P \leq 0.001$

**Figure S3**

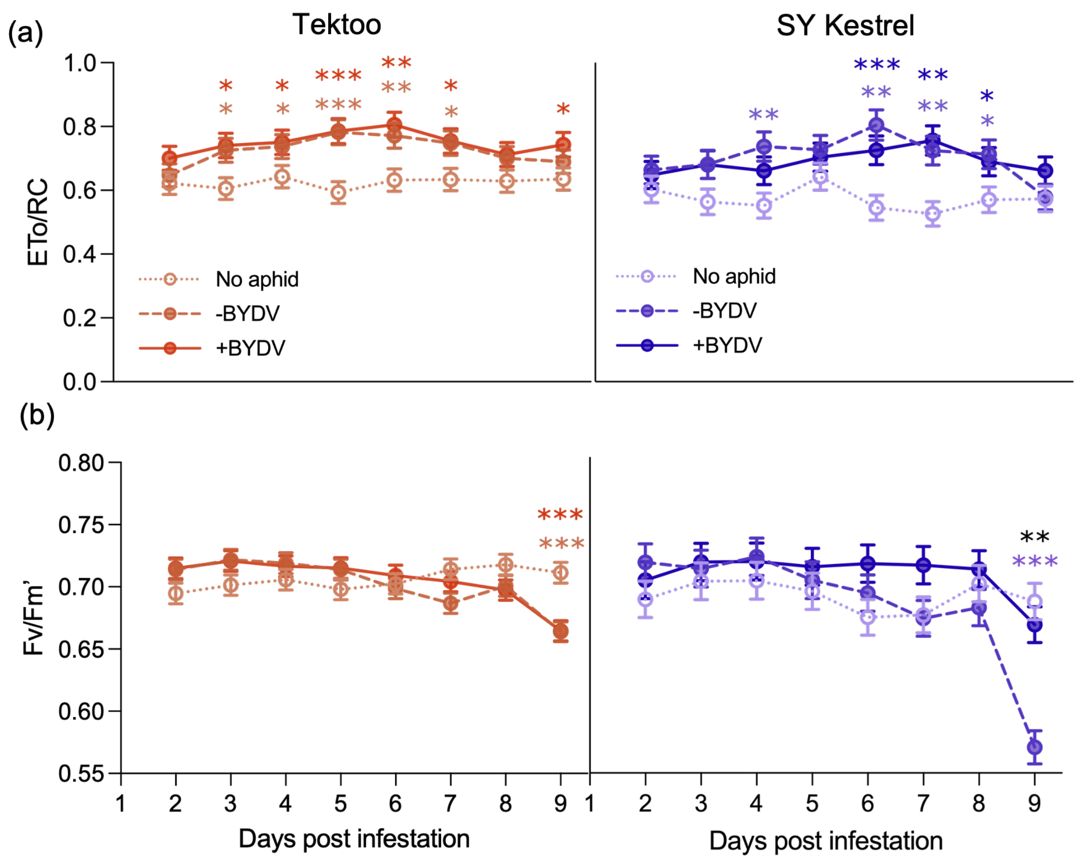


**Fig S3.** Chlorophyll fluorescence parameters in the barley hybrids SY Kestrel (Ryd4 introgression) and Tektoo (lacking the introgression) following infestation with non-viruliferous (-BYDV) or viruliferous (+BYDV) *R. padi*, or in uninfested controls, from 2 to 9 d post infestation (dpi) **(a)** Electron transport rate per reaction centre (ETo/RC) and **(b)** maximum potential quantum yield (Fv/Fm') of no aphid, -BYDV and +BYDV Tektoo and SY Kestrel from 2 to 9 days post infestation. All chlorophyll fluorescence parameters were analysed using a generalised linear model (GLM) with poisson distribution and log link function. ($n$ = 6). Back-transformed means shown. * $P \leq 0.05$; ** $P \leq 0.01$; *** $P \leq 0.001$ Stars denoting significant differences in black are differences between -BYDV and +BYDV treatment, stars denoting significant differences in colour of -BYDV or +BYDV treatment are in comparison to the no aphid treatment.

### Figure S4

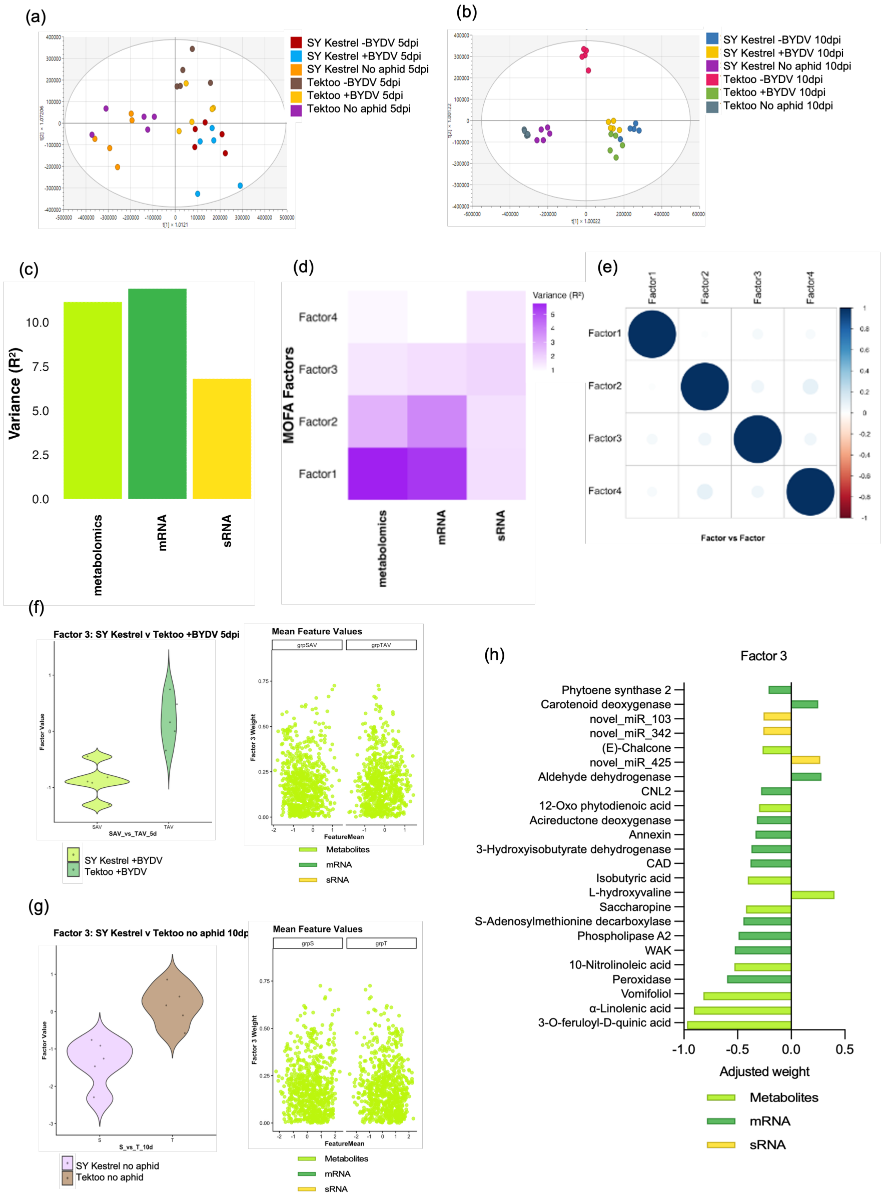


**Fig S4.** Orthogonal partial least squares-discrimination analysis (OPLS-DA) plot and multi-omics factorial analysis (MOFA)for the barley hybrids SY Kestrel (Ryd4 introgression) and Tektoo (lacking the introgression) following infestation with non-viruliferous (-BYDV) or viruliferous (+BYDV) *R. padi*, or in uninfested controls. OPLS-DA plot of barley metablic profile at (**a**) 5 days post infestation (dpi) and (**b**) 10dpi. Variance distribution **(c)** in overall dataset views and **(d)** per factor views. **(e)** Correlation between factors. Feature distribution and mean feature values in factor 3, separating **(f)** BYDV infected SY Kestrel from Tektoo treated at 5dpi and **(g)** no aphid treatted SY Kestrel from Tektoo at 10dpi. **(h)** Weights of the key features linked to group separation in MOFA factor 3.

### Figure S5

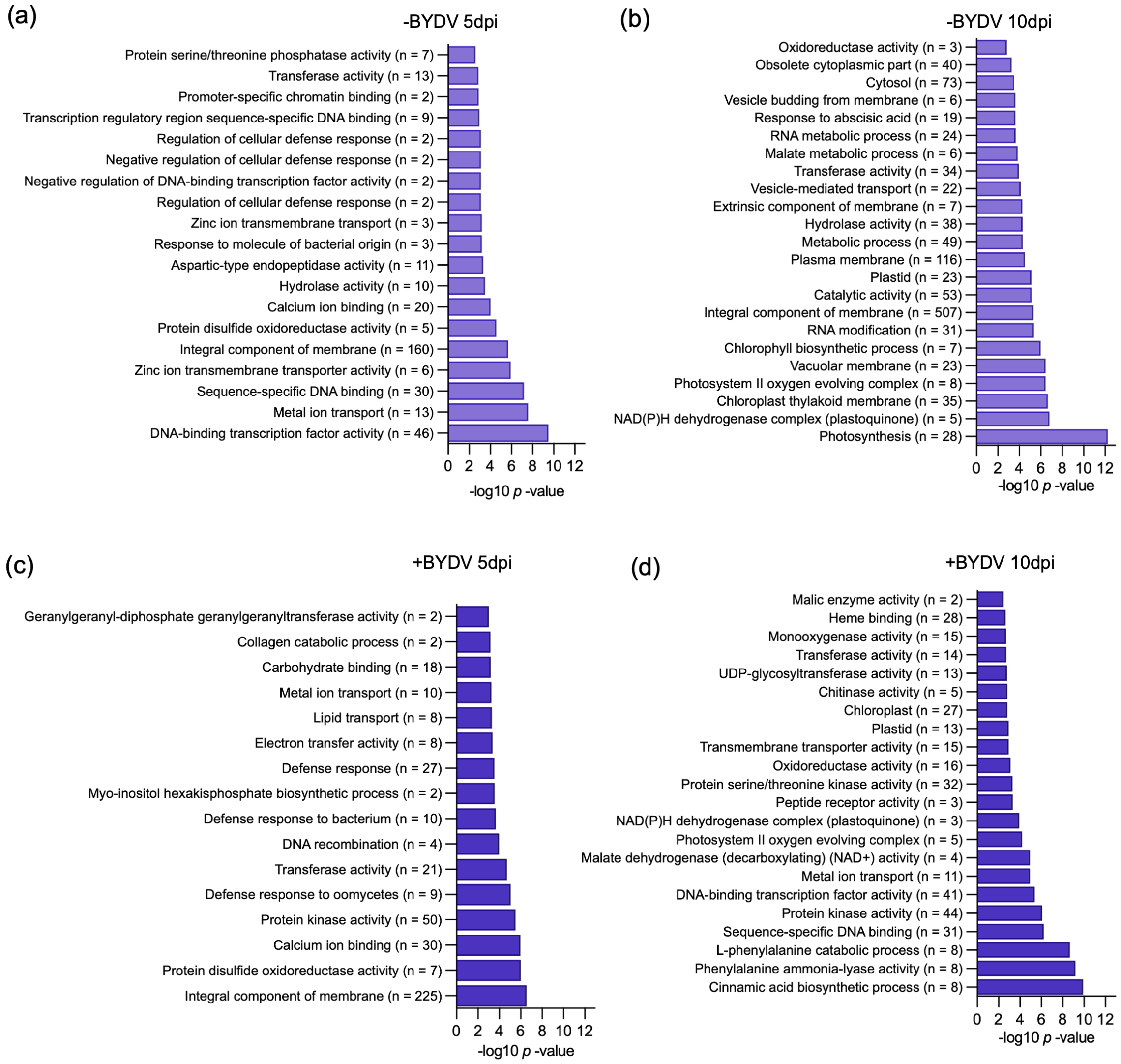


**Fig S5.** Gene Ontology (GO) enrichment analysis of the barley hybrids SY Kestrel (Ryd4 introgression) and Tektoo (lacking the introgression) subjected to no aphid, non- viruliferous (-BYDV) and viruliferous (+BYDV) *Rhopalosiphum padi* infestation. Differential gene expression of significant pathway terms with *p*-value <0.05 from samples under **(a)** –BYDV infestation harvested at 5days post infestation (dpi) **(b)** and 10dpi, **(c)** viruliferous (+BYDV) *R. padi* infestation harvested to 5dpi **(d)** and 10dpi.

**Figure S6**

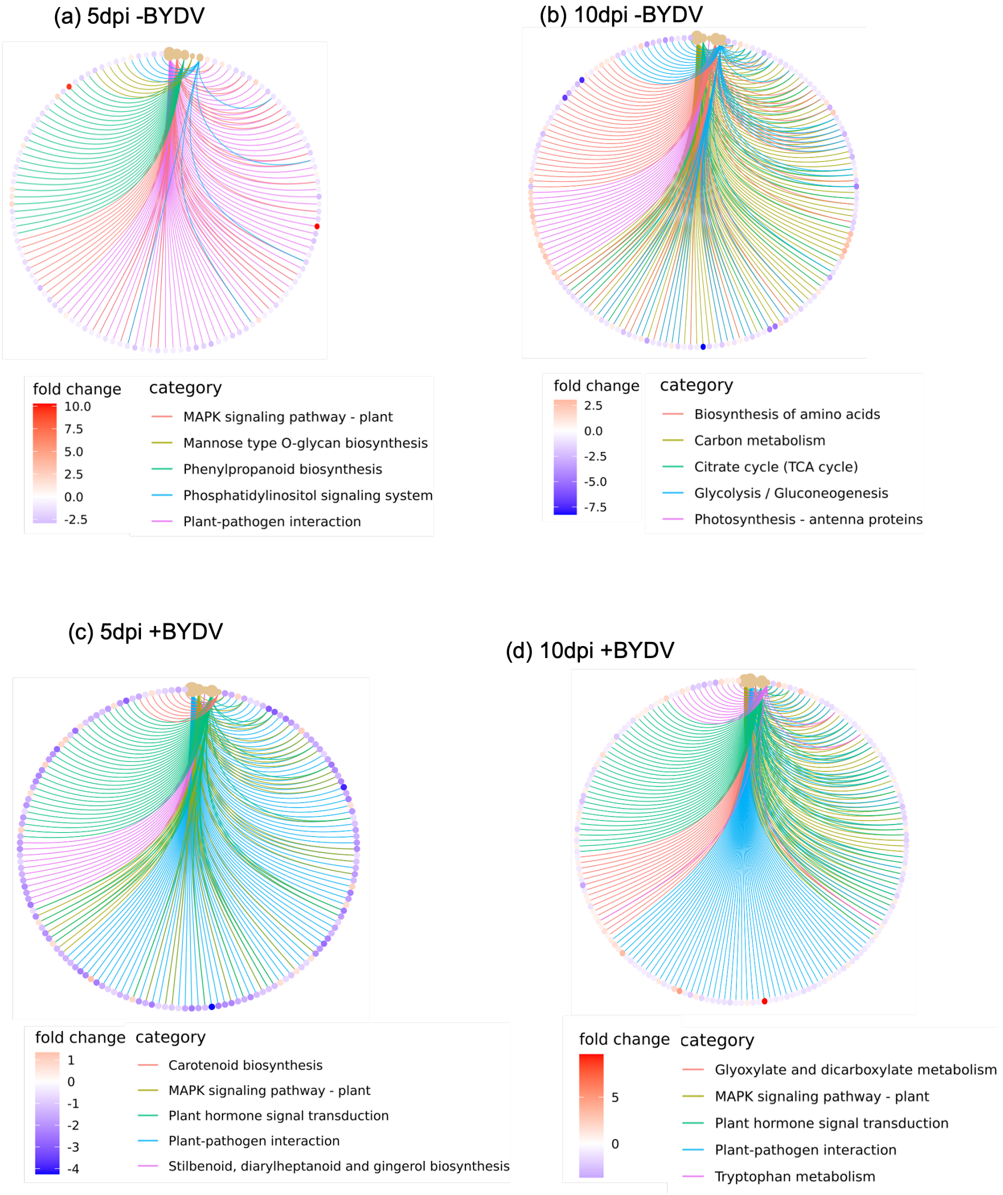


**Fig S6.** Kyoto Encyclopedia of Genes and Genomes (KEGG) pathway enrichment analysis of the barley hybrids SY Kestrel (Ryd4 introgression) and Tektoo (lacking the introgression) subjected to no aphid, non- viruliferous (-BYDV) and viruliferous (+BYDV) *Rhopalosiphum padi* infestation. Differential gene expression of significant pathway terms with *p*-value <0.05 from samples under **(a)** –BYDV infestation harvested at 5days post infestation (dpi) **(b)** and 10dpi, **(c)** viruliferous (+BYDV) *R. padi* infestation harvested to 5dpi **(d)** and 10dpi.
